# Visual cortical area contributions to the transient, multifocal and steady-state VEP: A forward model-informed analysis

**DOI:** 10.1101/2023.09.01.555874

**Authors:** Kieran S Mohr, Anna C Geuzebroek, Simon P Kelly

## Abstract

Central to our understanding of how visual evoked potentials (VEPs) contribute to visual processing is the question of where their anatomical sources are. Three well-established measures of low-level visual cortical activity are widely used: the first component (“C1”) of the transient and multifocal VEP, and the steady-state VEP (SSVEP). Although primary visual cortex (V1) activity has often been implicated in the generation of all three signals, their dominant sources remain uncertain due to the limited resolution and methodological heterogeneity of source modelling. Here, we provide the first characterisation of all three signals in one analytic framework centred on the ‘cruciform model’, which describes how scalp topographies of V1 activity vary with stimulus location due to the retinotopy and unique folding pattern of V1. We measured the transient C1, multifocal C1, and SSVEPs driven by an 18.75Hz and 7.5Hz flicker, and regressed them against forward-models of areas V1, V2 and V3 generated from the Benson-2014 retinotopy atlas. The topographic variations of all four VEP signals across the visual field were better captured by V1 models, explaining between 2-6 times more variance than V2/V3. Models with all three visual areas improved fit further, but complementary analyses of temporal dynamics across all three signals indicated that the bulk of extrastriate contributions occur considerably later than V1. Overall, our data support the use of peak C1 amplitude and SSVEPs to probe V1 activity, although the SSVEP contains stronger extrastriate contributions. Moreover, we provide elaborated heuristics to distinguish visual areas in VEP data based on signal lateralization as well as polarity inversion.

## 1 Introduction

Visual evoked potentials (VEPs) - electrical scalp potentials emanating from the brain in response to visual stimuli - are a central tool for electrophysiological work in both basic and clinical research domains to study the mechanisms of visual-cognitive processes and how they differ in a variety of clinical disorders (Creel, 2019; Kothari et al., 2016). Three main VEP-eliciting techniques are widely used to probe low-level visual processing in distinct, complementary ways. First, *transient VEPs* elicited by abruptly presented, isolated stimuli reveal a full cascade of processing stages in response to a discrete sensory event, with the earliest, “C1” component (60-90 ms) drawing particular interest for questions regarding how soon visual processing can be influenced by cognitive factors such as attention (Luck, 2014). Second, the *steady-state VEP* (SSVEP), a narrow-band spectral component generated by rhythmically flickering stimuli, can be used to trace evoked response amplitude continuously over time for a prolonged stimulus (Regan, 1966). Third, *multifocal VEPs* (mfVEP) are generated using rapid, statistically precise stimulation protocols across the full visual field (Baseler et al., 1994; James, 2003; see Lalor et al., 2006 for a related technique), providing a temporal profile of aggregate, rudimentary processing of pulses within a long, rapid sequence. While the transient VEP offers a view of more processing stages, mfVEPs and SSVEPs offer high signal-to-noise ratios (SNR) from shorter recording times, as well as the simultaneous probing of multiple visual locations.

Given the prominence and complementarity of these three distinct VEP signals, it is important to know what their cortical sources are, not only to expand the insights that they can provide individually but also to understand how they relate to one another. Source analysis studies have modelled the cortical origins of early visual components such as the C1 component of both the transient (Di Russo et al., 2002, 2005) and multifocal (James, 2003; Slotnick et al., 1999) VEP, as well as the SSVEP (Di Russo et al., 2007), with the implicated areas typically including primary visual/striate cortex (V1). However, much uncertainty remains regarding the dominant sources of these signals due to the ill-posed nature of the inverse problem in source analysis - that many dipole configurations can explain a given scalp potential distribution. This is particularly problematic for the visual system where many distinct areas lie in close proximity or have similar cortical orientation, which can lead to “crosstalk”, where signals from one area masquerade as those from another (Bradley et al., 2016; Dale & Sereno, 1993; Felleman & Essen, 1991; Grech et al., 2008; Hauk et al., 2022; Zorzos et al., 2021). What’s more, there exist many different source analysis methods, with different settings and assumptions to constrain them, which can yield different source estimates (Jonmohamadi et al., 2014; Mahjoory et al., 2017).

However, the fact that visual brain regions are retinotopically organised (Felleman & Essen, 1991) can provide crucial additional traction on the problem of distinguishing their scalp-projected activity. This is particularly true at early stages of the visual hierarchy where receptive fields are small (Dumoulin & Wandell, 2008), allowing characteristic cortical positions and folding patterns to make clear predictions of retinotopically varying scalp topographies. Indeed, this forms the basis of recent, improved source analysis methods that constrain estimates using functional retinotopic maps (J. Ales et al., 2010; Hagler et al., 2009; Hagler, 2014; Hagler Jr. & Dale, 2013). The primary example of this kind of geometric means of source identification is the “cruciform model” of V1 morphology. V1 extends over a medial-occipital area encompassing the Calcarine sulcus, with the upper visual field projecting to its ventral floor and lower field to its dorsal ceiling, and the medially-facing banks of the sulcus where visual field locations near the vertical meridian are represented (Benson et al., 2014; Halliday & Michael, 1970; Wandell et al., 2009). This geometry predicts that a V1 signal should invert in polarity when stimuli appear in the upper versus the lower visual field and become increasingly lateralized as stimuli are presented closer to the vertical meridian. This logical principle formed the basis of the original claims that the C1 component of the transient VEP is generated in V1 (Clark et al., 1994; Jeffreys & Axford, 1972). The same logic has also been applied to the C1 component of the mfVEP (Lalor et al., 2012; Slotnick et al., 1999), but not yet to the SSVEP, the sources of which have thus far been indicated only by source analysis (Di Russo et al., 2007).

The validity of relying on the cruciform model to identify V1 as a source for scalp potentials has, however, been debated (Ales et al., 2013; Kelly, Schroeder, et al., 2013; Kelly, Vanegas, et al., 2013). Some studies had focussed on an abbreviated form of the cruciform model centred on polarity reversal across the horizontal meridian, but a problem with this is that extrastriate areas V2 and V3 also reverse in cortical surface orientation between the upper and lower visual field (Edwards & Drasdo, 1987; Halliday & Michael, 1970; Maier et al., 1987). Ales et al (J. M. Ales et al., 2010) corroborated this with fMRI-informed topographic simulations, arguing therefore that polarity inversion across the horizontal meridian is not a unique identifier of V1. Ales et al (Ales et al., 2013) further pointed out that some animal neurophysiology work indicates considerable overlap in the latencies of responses in V1 and many extrastriate areas (Bullier & Nowak, 1995; Chen et al., 2007; Maunsell & Gibson, 1992; Nowak et al., 1995; Raiguel et al., 1989; Robinson & Rugg, 1988; Schroeder et al., 1998). These points together imply that early, polarity-inverting VEP components like the C1 might be as consistent with V2/V3 as they are with V1. However, while this indeed undermines the sole use of polarity reversal to identify a V1 source (J. M. Ales et al., 2010), it fails to take account of additional topographic shifts predicted by the full cruciform model for stimuli near the vertical meridian, where foci should become more ipsilateral near the upper vertical meridian and more contralateral near the lower vertical meridian (Kelly, Vanegas, et al., 2013). Since V2 and V3 do not mimic this aspect of V1 geometry, a full consideration of the cruciform model across the entire visual field should generate topographic predictions unique to V1.

The goal of this study was to carry out a full “cruciform analysis” of the transient VEP C1, the mfVEP C1 and the SSVEP to establish the extent to which retinotopically varying topographies can distinguish between sources in V1 and V2/V3 due to their distinct cortical morphology. Further, we leverage all three signals to estimate response latency differences across the three visual areas. To do this, we recorded transient VEPs, mfVEPs, and SSVEPs at fast and slow flicker rates, and analysed their topographic variation across the visual field with respect to forward-model predictions derived from the publicly available Benson-2014 retinotopy atlas (Benson et al., 2014) for V1, V2 and V3. This study complements recent work using retinotopy-constrained source estimation (RCSE) methods to distinguish V1, V2 and V3 activity (J. Ales et al., 2010; Hagler, 2014; Hagler et al., 2009; Hagler Jr. & Dale, 2013) by taking a distinct viewpoint and approach. Rather than using individual functional retinotopic maps to fit individual VEP data, we used grand-average retinotopic maps to fit grand-average VEP data. There is an inherent trade-off between these approaches from a source-modelling quality perspective. Using individual data unsmoothed by averaging offers potentially more distinctive topographic variation across locations and areas to constrain the models and distinguish those areas, but reaping this benefit depends on how accurately those individual variations are measured despite noise in individual fMRI, which in practice usually calls for additional model adjustments to improve fit. Ultimately, modelling at the grand-average level allowed us to focus on the characteristic topography features across the visual field for sources in different visual areas, allowing us to provide a pathway to understand the VEP topographical trends indicative of different visual areas and thus enable a degree of reliable source inference in the absence of individual fMRI data. In so doing, we established that V1 makes the dominant contribution to all four VEP signals. Although extrastriate areas were able to explain some variance both on their own and alongside V1, when included with V1 in the same regression models of the full visual field, their weights did not build substantively until distinctly beyond the C1 peak in the transient VEP, and built little if at all in the mfVEP. Meanwhile, modelling full sinusoidal time courses for the fundamental SSVEP frequencies revealed frequency-dependent relative extrastriate contributions and possible response lags for V2 and V3 on the order of tens of milliseconds, relative to V1. Finally, our topographic analyses suggest an update to the heuristic used to distinguish V1 sources from V2/V3 sources given VEPs from as few as one upper- and one lower-field location: alongside the classic polarity reversal between upper and lower visual field locations, the dominant focus of upper field V1 topographies tend to be ipsilateral while lower field ones tend to be contralateral.

## 2 Method

### 2.1 Participants

Ten healthy young adults took part in this experiment (5 Females, 10 right handed) with a mean age of 23.5 years (SD=3.3). They were compensated for their participation with a lump sum of €20. All participants gave written informed consent to be included in the study, were over the age of 18, had normal or corrected to normal vision and reported no neurological or psychiatric conditions. All operations were approved by the Human Research Ethics for Sciences board of University College Dublin and adhered to the guidelines set out in the Declaration of Helsinki.

### 2.2 Stimuli

Stimuli were presented using Psychtoolbox-3 (Brainard, 1997; Kleiner et al., 2007; Pelli, 1997) in a dark, sound-attenuated chamber on a 1024 × 768 Dell E771p CRT monitor (32.5 × 24.5cm) operating at a refresh rate of 75Hz. Participants were seated at a chin rest at a distance of 57 cm from the monitor. They maintained a stable gaze on a white fixation cross, which was displayed against a uniform grey field (64cd/m^2^, half the monitor’s maximum luminance). Stimuli were constructed from an annular “dartboard” pattern of alternating black and white checks, and had an inner and outer radius of 2.75° and 7.25° (see Figure 1A). This was divided angularly into 16 wedges of 22.5° polar angle, each containing a 4 × 4 check pattern (angle × eccentricity, equally spaced), which spanned the visual field. This full-field stimulation allowed us to measure the retinotopic patterns of visually evoked EEG topographies and compare these to predictions made by the Benson-2014 retinotopic atlas (Benson et al., 2014). The retinotopic layout of V1 and V2 in this average brain is shown in Figure 1B-C, demonstrating the cruciform morphology of V1 as it passes through the Calcarine sulcus. This morphology motivates the cruciform model of V1 (Fig. 1D) as a means of determining striate origin for scalp signals, which should exhibit a specific sequence of topography shifts as a function of retinotopic stimulus position. Figure 1D outlines the cortical morphology of V2 and V3 along with V1, all of which predict polarity reversal across the horizontal meridian. Crucially however, V1 predicts a pattern of topography shifts within quadrants of the visual field that is not shared by V2 or V3: ipsilateral shifts between the horizontal meridian and the upper vertical meridian, and contralateral shifts between the horizontal meridian and the lower vertical meridian.

**Figure 1:**
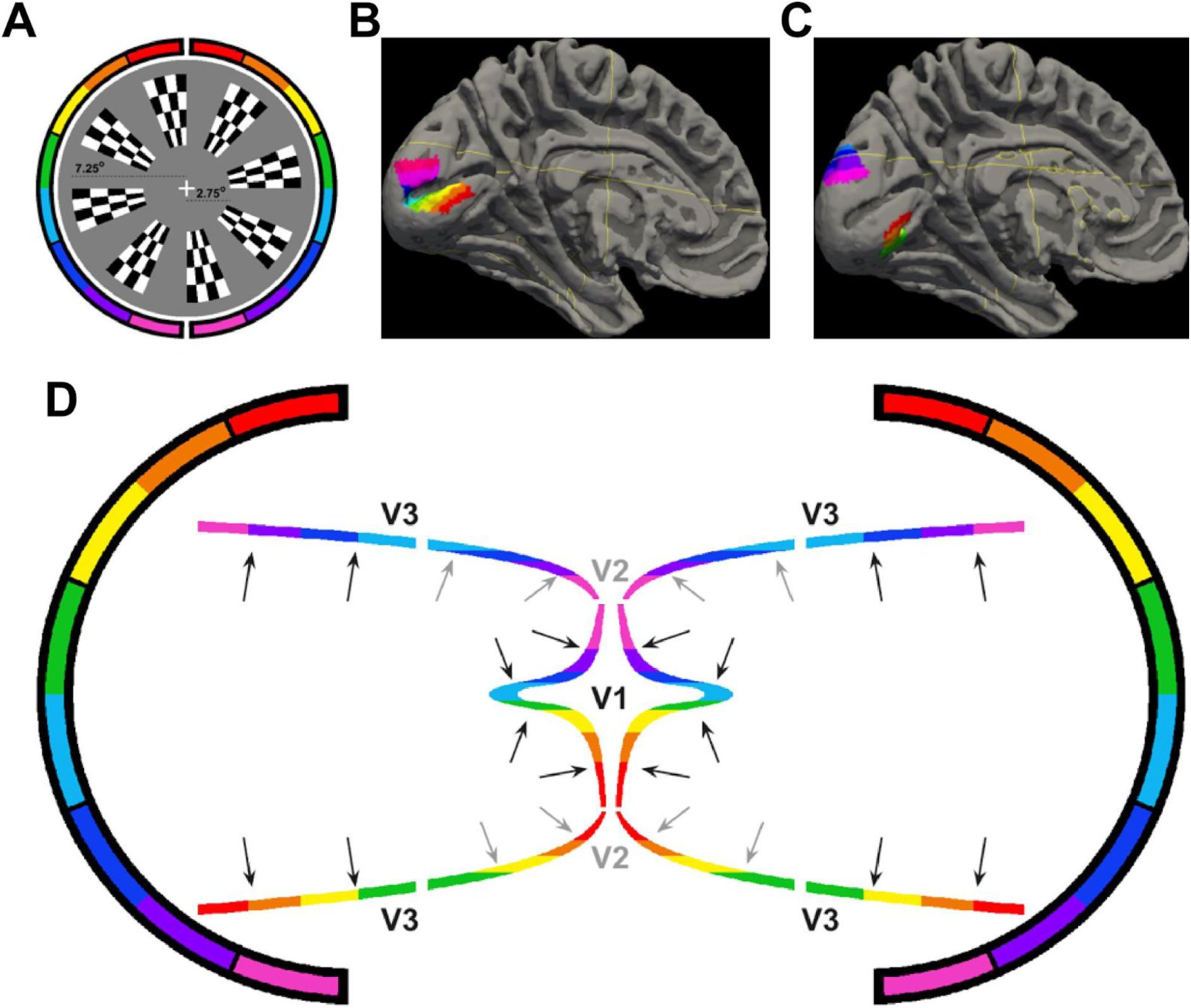
**A)** Stimuli used across all visual stimulation protocols. For illustration, every other wedge is displayed (there were 16 in total) with the inner and outer eccentricity indicated by the lengths of the two dashed lines. **B-C)** Left hemisphere of the MNI-152 brain fitted with the Benson-2014 atlas, showing the retinotopic map positions of the 8 checkerboard wedges in the right visual field for V1 (B) and V2 (C). **D)** Illustration of the cruciform model. A cartoon approximation of the cortical folding pattern is shown for V1, V2 and V3 as viewed from a coronal slice. The outer brackets represent regions of the visual field from the lower vertical meridian (pink/purple) to the upper vertical meridian (red/orange) illustrating the up-down image reversal between visual images and cortical retinotopy (the left-right image reversal is not colour-coded in this cartoon). The grey arrows depict examples of surface normal dipoles and outline the different within-quadrant dipole shifts expected of each visual area.

### 2.3 Procedure

Participants observed a series of visual stimulation protocols designed to elicit a transient VEP, a pattern-pulse mfVEP, and an SSVEP, for each of the 16 wedges. They were given no task other than to maintain steady eye gaze on a central fixation cross. Across the three protocols, overall stimulation time was approximately 70 minutes and participants had the opportunity to take regular breaks.

The transient VEP protocol consisted of 200 trials for each location in which stimuli were flashed for 50 ms in random sequence, separated by 500 ms. This was divided into seven blocks of approximately seven minutes each. In the mfVEP protocol, the 16 wedges followed orthogonal rapid pulse sequences that were governed by 16 time-lagged copies of a zero-autocorrelation series of binary numbers derived from a 4095 frame m-sequence (Baseler et al., 1994; James, 2003). Stimulus pulses lasted two frames (26.6 ms) and occurred at transitions from −1 (the “off digit”) to 1 (the “on digit”). To limit the overall pulse speed, three out of every four pulses were discarded, as has been done elsewhere (Vanegas et al., 2013), which yielded a protocol with 256 stimulus pulses for each location extending across a time period of 54.6 seconds. This was repeated 10 times with different m-sequences yielding 2,560 pulses per location. The SSVEP protocol included a fast (18.75 Hz) and a slow (7.5 Hz) subdivision. The 18.75 Hz flicker was produced by a sequence of two “on” and two “off” frames. The 7.5 Hz flicker was produced by a sequence of ten frames (5 on and 5 off). Each wedge flickered at both frequencies for six unbroken periods of 5 seconds. To reduce overall stimulation time, two locations 180° apart flickered simultaneously, one at each frequency. Locations were stimulated in random order, separated by a self-paced gap (the participant clicked the mouse to proceed to the next location).

### 2.4 Data Acquisition

EEG data were recorded at 512 Hz by an ActiveTwo Biosemi system with 128 scalp electrodes following the Biosemi ABC layout (Biosemi, The Netherlands) and six external flat-faced electrodes placed above and below the left eye, on the left and right outer canthi, and on the left and right mastoids. Eye gaze and blinks were monitored via the four electrooculograms and using an Eyelink Plus 1000 Tower system (SR Research, ON, Canada) recording at 1000 Hz.

### 2.5 Data Processing

EEG processing was carried out using a combination of inhouse Matlab scripts (Mathworks, The United Kingdom) and EEGLAB routines (Delorme & Makeig, 2004). These processes were similar across the three stimulation protocols except that trials containing artifacts were not rejected for the mfVEP or the SSVEP as this would complicate the analysis due to their continuous stimulation streams. This is justified since both signals are robust to artifacts because the SSVEP has a much narrower spectral band than do artifacts and the mfVEP has a very large number of trials making it robust to occasional artifacts.

Across the board, continuous data were low pass filtered by convolution with a 77-tap hanning-windowed sinc function with a 3-dB corner frequency of 35.3 Hz and 50 Hz attenuation (mains) of 83.5 dB. Long segments of persistently high noise in individual channels were linearly interpolated using EEGLAB’s *eeg_interp* function. To identify these, the data were partitioned into short segments of 5 seconds and low- and high-frequency standard deviation (following a 5 Hz low pass filter and subtraction of this filtered data, respectively) were calculated and subjected to thresholds of 20 mV and 10 mV. Only long segments with a consistent sequence of 7 or more such segments were interpolated and short gaps between such long segments (2 or fewer segments) were also interpolated.

The data were then re-referenced to the average of all scalp channels and low frequency trends were removed. To avoid distortions that can result from typical high pass filters (Acunzo et al., 2012), a linear detrending approach was taken instead. This entailed partitioning the data into segments and detrending each segment by subtracting the line of best fit for each electrode. The segment durations were adjusted slightly so that the partition edges did not coincide with any epoch window. These durations differed across the stimulation protocols to take their individual requirements into account. For the transient VEP, a duration of 4 seconds was used in order to span multiple trials and avoid biassing the fitted line with any trial order effects. For the SSVEP, where unbroken epochs of 4 seconds were needed, the duration was set to 5 seconds. Finally, for the mfVEP, where at any moment in time a large number of trials are at various stages of their time course and avoiding partition points inside epochs is impossible, a shorter duration of 500ms was used.

At this point, for the transient VEP, continuous data were then subjected to in-house artifact detection routines to identify and label time points with electrode “pops”, slow wave drift, muscle activity and blinks. Pops were identified by the absolute value of a 20-ms-lag amplitude difference exceeding 50 mV. Slow drift and muscle activity artifacts were detected using discrete Fourier transforms of successive one-second partitions. Slow drift was identified if a ratio exceeding 5:1 was found between the maximal frequency-domain amplitudes in the 0-3 Hz and the 3-7 Hz ranges, and muscle activity was identified if a similar ratio between the 20-40 Hz and 3-7 Hz ranges exceeded 2:1. Blinks were identified both with the eye-tracking data (for full blinks) and via the difference between the upper and lower vertical electrooculograms (for partial blinks), which was transformed by taking its first derivative (to remove any slow baseline shifts) and applying a 200 ms moving average (to extend blink detection to the periods immediately before and after the main blink where EEG may still be affected). Data points where this signal exceeded 1 mV were labelled as blinks. Eye tracking data was used to identify saccades, which were defined as gaze displacements over a 10 ms period in excess of 0.2°. Average trial-loss rate across participants was 11.3%, with a maximum of 28.7% (leaving a minimum of 142 trials per participant per location).

### 2.6 VEP signal derivation

For the transient VEP and the mfVEP, data were divided into stimulus-locked epochs ranging from −100 ms to 400 ms and averaged for each of the 16 wedges separately. Note that for the mfVEP, the resultant waveforms were almost identical (*r=0.98*) when the continuous data were regressed against the stimulus pulse matrix. Grand average waveforms for each signal and wedge were then produced by averaging across participants. Topographies were calculated between 70 and 80 ms to correspond with the C1 timeframe.

For the SSVEP, the first 500 ms after flicker onset were discarded in order to exclude the onset-related VEP, and a four-second epoch was extracted from that point onwards. Each of these four-second epochs (six per location and SSVEP frequency) were transformed to the frequency domain by means of a discrete Fourier transform and the flicker frequencies of 18.75 Hz and 7.5 Hz were extracted from the spectrograms. These were then averaged across repetitions to yield participant averages, which were in turn averaged across participants. To capture polarity and produce topographies similar to the C1 above, we reasoned that SSVEP oscillations from a given visual region would have similar phase across electrodes and visual field locations (Hagler, 2014) but be phase-shifted by 180° at electrodes where polarity was reversed. We thus determined the “principal phase axis” - the line through the origin in the complex plane to which all electrodes and retinotopic locations best align - and projected each complex FFT value onto this axis. We assigned positive and negative polarity to the two ends of the principal phase axis in such a way that upper field locations close to the horizontal meridian produced negative topographic foci in accordance with the C1. Importantly, because this choice does not affect the within-quadrant topography shifts that are differentially characteristic of V1, V2 and V3, it does not bias the results to demonstrate striate generation of the SSVEP or topographic similarity between the SSVEP and the C1.

### 2.7 Forward Modelling

To predict EEG topographies for sources in V1, V2 and V3 based on the stimuli used to elicit the empirical VEPs, the cortical surface area corresponding to each wedge was estimated in an MNI-ICBM152 non-linear average brain (Grabner et al., 2006). To do this, the Benson-2014 retinotopic atlas (Benson et al., 2014) was fit to the MNI-152 average brain, which facilitated the estimation of retinotopic maps for the visual areas. Following reconstruction and volumetric segmentation of the MNI-152 average brain using Freesurfer (http://surfer.nmr.mgh.harvard.edu/; Dale et al., 1999; Fischl et al., 1999; Fischl & Dale, 2000; Salat et al., 2004), the Benson-2014 atlas was applied using the *Neuropythy* Python module (Benson & Winawer, 2018) and regions of interest (ROIs) were defined for the segments of the visual field encapsulated by each of the 16 wedges for each visual area. This was used to generate a list of voxel coordinates corresponding to the grey matter surface area for each wedge in each visual area.

The Matlab toolbox Fieldtrip (Oostenveld et al., 2010) was then used to generate a conductivity head-model based on this brain (boundary element model; BEM). To generate a forward-model, this was co-registered with the Biosemi electrode coordinates by manually rotating the electrode coordinates to align with the head model. The forward-model was then used to derive scalp topographies resulting from dipoles located at each of the voxels determined by the above ROIs, the orientations of which were constrained by the surface normal of the corresponding cortical surface area. To find the surface normal at a given voxel, we applied linear regression to the 5×5×5 cube of voxels (5 mm^3^) centred on it to find the line that best predicted voxel brightness values from their position coordinates, or in other words, best discriminated white from grey matter. These single-voxel topographies were then averaged over all voxels corresponding to each wedge (assuming uniform activation across all voxels) to yield the final predicted topographies for each wedge. This whole process was repeated for V1, V2 and V3 to yield predictions of retinotopically mapped topographies for each wedge and each of the 3 visual areas.

## 3 Results

### 3.1 V1 provides the best fit to empirical VEPs

Figure 2 shows the grand average topographies of each empirical VEP signal (panels A-D) and the topographies predicted for V1-3 based on the Benson-2014 retinotopy atlas (panels E-F). Each of the VEPs demonstrated a pattern of topographies that inverted in polarity between the upper and lower visual field. In addition to this, they tended to be ipsilateral in the upper visual field and contralateral in the lower field, shifting laterally in these same directions as stimuli came closer to the vertical meridian within each quadrant. The only exception to this trend was the lower field of the 7.5Hz SSVEP, which showed a surprising pattern that we will address later (see section “Phase shift in the 7.5Hz SSVEP”).

**Figure 2:**
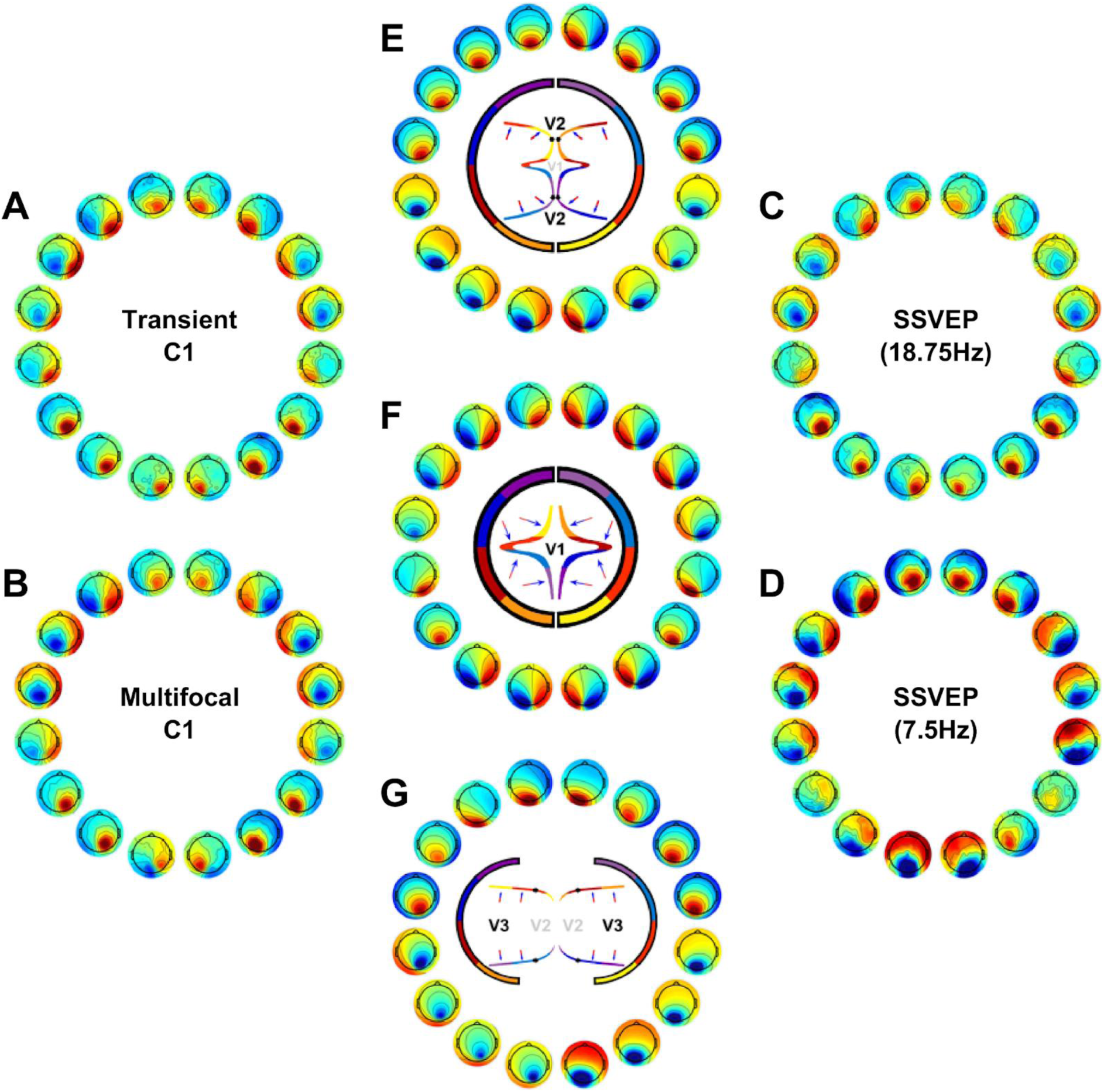
**A-D)** Grand average topographies at the 16 stimulated locations for the transient C1 (A), the multifocal C1 (B), the 18.75 Hz SSVEP (C), and the 7.5 Hz SSVEP (D). **E-G.** Topographies predicted for V1 (F), V2 (E) and V3 (G) based on retinotopic data from the Benson-2014 atlas fit to the MNI-152 brain. Inside the circle of topographies is a cartoon depiction of the cruciform model. The outer brackets provide a colour-coding scheme of visual field octants such that the inner depiction of cortical morphology shows an approximation of the retinotopic map layout. Black markers highlight the transition between V1 and V2 (E), and between V2 and V3 (G). The blue and red arrows demonstrate typical dipole directions for each octant with the head of the arrow representing the negative end, under the assumption of surface-negative potentials.

The MRI-predicted topographies for V1 had a similar pattern, also showing polarity inversion across the horizontal meridian, ipsilateral upper-field topographies and contralateral lower-field topographies, as well as within-quadrant patterns of upper-ipsilateral and lower-contralateral shifts (Fig. 2F). This is because in V1, locations near the vertical meridian are represented along the medial wall whereas those near the horizontal meridian are represented inside the Calcarine sulcus (Fig. 2F, also see Fig. 1B). V2 (Fig. 2E) and V3 (Fig. 2G) predicted polarity inversion across the horizontal meridian as well (in this case positive for upper and negative for lower field) but they tended to predict contralateral topographies in the upper field and V2 topographies tended to be ipsilateral in the lower visual field. Within quadrants, V2 topographies shifted in the opposite direction to those of V1 and for V3 they shifted little if at all.

To measure these qualitative patterns quantitatively, correlations (Pearson’s *r*) were calculated between the empirical and MRI-predicted topographies (treating electrodes as observations). To capture changes in similarity across visual field locations, these correlations were calculated for each location separately (Fig. 3). The qualitative patterns outlined above were largely recapitulated. V1 showed positive correlations at all locations for all VEPs except for the 7.5 Hz SSVEP in the lower field; V2 showed negative correlations near the horizontal meridian and positive correlations near the vertical meridian reflecting its prediction of an opposite pattern of within-quadrant topography shifts; V3 showed negative correlations at most, but not all, locations.

**Figure 3:**
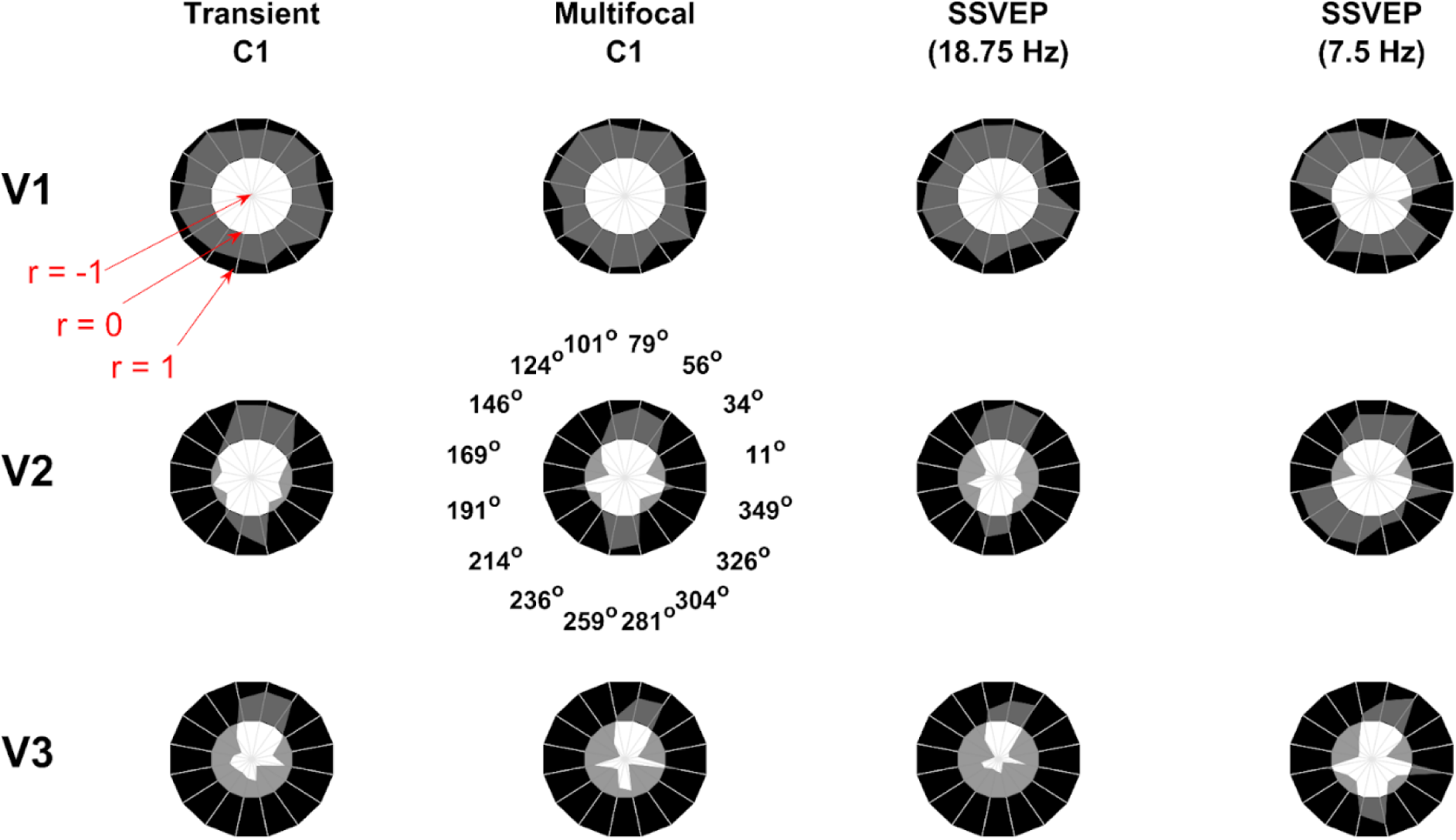
Correlations between MRI-predicted topographies for V1-3 and each of the VEP signals as a function of stimulus location. The angles shown for V2 and the multifocal C1 apply to all pairings. In the annular plots, the outer black radius corresponds to a correlation of *r=1*, the interface between the black and white sections corresponds to *r=0*, and the origin (centre of the white inner circle) corresponds to *r=-1*. Each ‘spoke’ of these annular plots aligns with the midpoint of the polar angle of each checkerboard wedge. Correlations are uniformly positive for V1 with the exception of the lower field for the 7.5 Hz SSVEP. Correlations for V2 are largely positive near the vertical meridian and negative near the horizontal meridian. Finally, correlations are largely negative for V3 except for the upper vertical meridian.

In order to quantify how well the predicted topographies captured the overall pattern of retinotopic topography shifts in the empirical VEP signals, topographic correlations were also calculated across all locations together by stacking observations into a single 2,048 long vector (128 electrodes × 16 locations). To assess statistical significance for these correlations, location was shuffled for the predicted topographies (within each visual area) and the correlations recalculated 1,000,000 times to generate null distributions. These distributions, along with their corresponding point estimates and p-values, are shown in Figure 4A. V1 significantly correlated with all four VEPs positively, whereas V2 correlated only with the 7.5 Hz SSVEP (positively), and V3 correlated negatively with all VEPs except the 7.5 Hz SSVEP.

**Figure 4:**
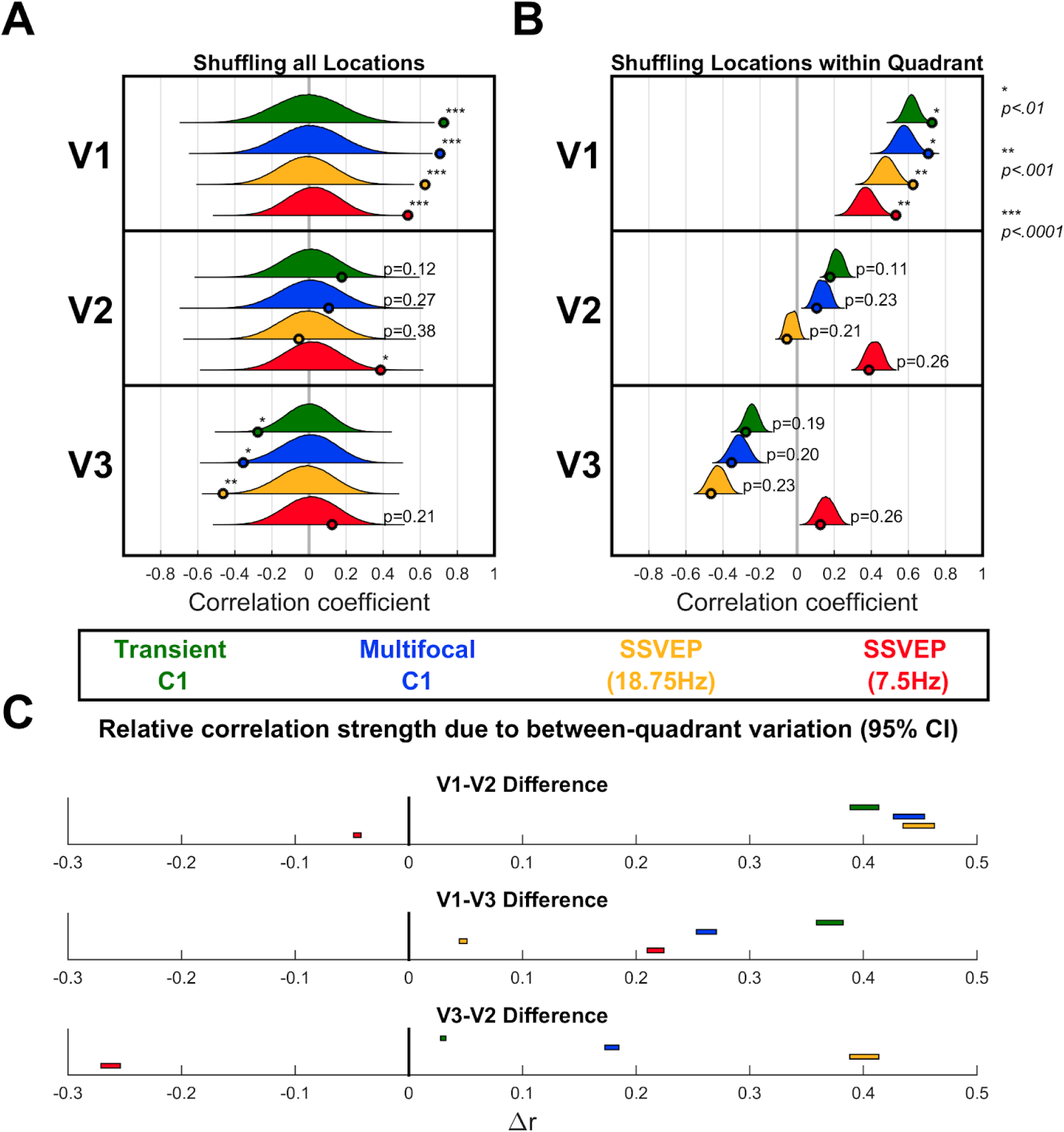
**A)** Correlations between MRI-predicted topographies for V1-3 and each of the VEP signals taking all locations together. The observed correlations (point estimates) are shown as small circles along the x-axis for each MRI-predicted topography (vertically stacked panels) and each VEP signal (colour-coded). In order to assess statistical significance, 1,000,000 random permutations of stimulus locations for the predicted topographies were taken and the correlations recalculated to generate a null distribution (shown alongside each point estimate). P-values corresponding to the observed point estimates are shown alongside them, with asterisks representing the following ranges: * *p<.01*, ** *p<.001*, *** *p<.*0001. **B)** In order to assess whether each visual area captured within-quadrant topography variations for each signal, these permutations were repeated, but only shuffling locations within each quadrant. These distributions are narrower and closer to their corresponding observed point estimates since they are constrained to retain predictive information stemming from between-quadrant topographic variations (e.g. polarity inversion between the upper and lower visual field). **C)** Since the null distributions in panel B retain between-quadrant variations, they serve to capture how well each visual area predicts the between-quadrant variations of each VEP signal. Here, the differences in the means of these distributions are calculated between each pair of visual areas for each visual component and shown as 95% confidence intervals (CIs). Where distributions were centred on negative correlations they were multiplied by −1 so that the CIs represented differences in absolute correlation.

To specifically assess whether the within-quadrant topographic variations were captured by each MRI-prediction, new null distributions were generated by shuffling locations only within their respective quadrants so that correlations that relied solely on between-quadrant topographic variations would no longer be significant. These distributions and the p-values for their corresponding point estimates are shown in Figure 4B. V1 continued to show significant positive correlations across the board while neither V2 nor V3 showed correlations that significantly differed from the new null distributions. These within-quadrant-shuffled null distributions themselves served a second purpose. They captured the component of topographic correlations due purely to between-quadrant topographic variations of the VEP signals. These between-quadrant variations are the most relevant for typical perceptual/cognitive VEP studies in which it is important to infer whether VEP signals originate in V1, because most studies do not fully map retinotopy and only have a single stimulus location in a given quadrant. As such, we compared these distributions for V1, V2 and V3, using permutation tests to generate confidence intervals of the pair-wise differences between the means (Figure 4C), to determine which visual area best accounts for the between-quadrant topographic variations in the VEP signals. These indicated that V1 provided a better match than V3, which in turn provided a better match than V2, in all cases (*p<.0001*) except the 7.5 Hz SSVEP for which V2 provided a better match than V1, which in turn provided a better match than V3 (*p<.0001*).

Since the above analysis demonstrated that V1 captures the between-quadrant topographic variation of most VEP signals better than V2/V3, we sought to identify the characteristic topographic features of each visual area and VEP signal that underlie this effect. As can be seen in Figure 2, the most prominent topographic features that varied across visual field locations were polarity and left-right lateralisation. However, the lateralisation of some topographies is difficult to appreciate visually. Therefore, we calculated a metric of left-right lateralisation to facilitate its comparison across VEP signals, visual areas and visual field locations. To do this, we first established a field-to-polarity mapping for each set of 16 topographies - i.e. whether the dominant topographic focus is negative among the upper field locations and positive in the lower field (as for the empirical C1), or vice versa. We then derived a centre-of-gravity measure of laterality for each individual location by taking a weighted average of the horizontal positions of all electrodes with that dominant polarity, with weighting proportional to signal amplitude. We multiplied by −1 as necessary so that positive values indicated a more ipsilateral location of the dominant focus in a given quadrant, whether positive or negative, and negative values indicated contralateral. This is plotted as a radial distance in Figure 5, showing that topographic foci for all VEP signals were ipsilateral in the upper visual field and contralateral in the lower visual field. This was also true of V1 topographies whereas the reverse was true of V2 topographies, and V3 topographies were closer to midline. This offers a valuable extension to heuristics that are applied to identify a VEP signal or component as having a source in V1 as opposed to extrastriate areas such as V2/V3 in studies that do not measure VEPs from the entire visual field: the dominant topographic focus should not only reverse in polarity between upper and lower-field locations but also reverse in laterality in this specific way. While this extended heuristic remains limited for locations very close to the vertical meridian, where V1 and V2 sources in particular are highly similar, it strengthens the case for locations away from the vertical meridian and accords with observed characteristics of the empirical C1 component (Di Russo et al., 2002).

**Figure 5:**
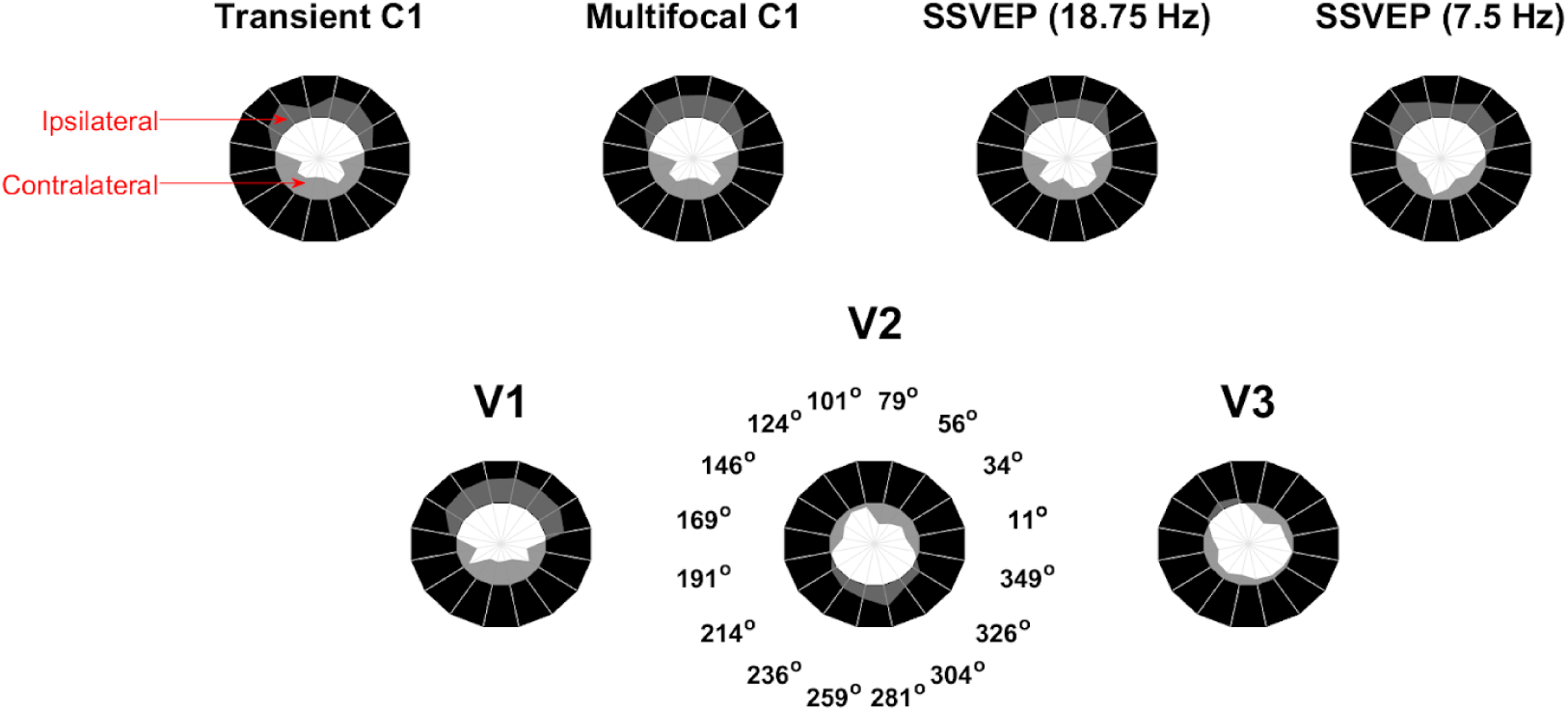
Measures of contralateral/ipsilateral topography locus as a function of visual field location for each of the VEP signals (top) and each visual area prediction (bottom). Radial distances further into the black annulus correspond to ipsilateral loci (with a limit of +1 reflecting the theoretical limit of all amplitude loaded on the most lateral electrode position) while radial distances further into the inner white circle correspond to contralateral loci (with a limit of −1, similarly for the other side). These lateralization values correspond to the negative part of topographies for upper visual field locations and the positive part for lower visual field locations, except for V2 and V3 simulations where the reverse mapping was used, in accordance with the polarity of the dominant foci in the upper versus lower field in each case. VEP signal topographies were ipsilateral in the upper visual field and contralateral in the lower visual field. V1 topographies were also ipsilateral in the upper visual field and contralateral in the lower visual field while the reverse was true of V2. V3 topographies were mostly close to the midline.

### 3.2 Modelling signal contributions from V1, V2 and V3

Individually, V1 provided a more complete account of the pattern of VEP signal topographies observed around the visual field than did V2/V3. However, this does not rule out the possibility that all three visual areas contribute to some degree to generating these signals. To address the extent of such contributions to each signal, we modelled various combinations of visual areas using multiple linear regression. Stacking all 16 visual field locations, each regression model had 2,048 observations (16 × 128) and the outcome variable was signal amplitude at each observed channel. The predictor variables were the MRI-predicted amplitudes for each visual area. All variables were converted to z-scores prior to carrying out the regressions. Seven such models were carried out for each VEP signal, one for each subset of the three visual areas: three models based on each visual area alone, three models based on each combination of two areas, and the model including all three visual areas.

The results of these models are shown in Figure 6, with details provided in Table 1. Among the single-area models, V1 explained the most variance (in line with the earlier analysis), explaining 53% for the transient C1, 50% for the multifocal C1, 39% for the 18.75 Hz SSVEP, and 28% for the 7.5 Hz SSVEP (compared with 8%, 13%, 22% and 15% for the next best fitting visual area in each case). Including all three visual areas explained additional variance (Figure 6A) and produced lower BIC values across the board (Figure 6B). This resulted in an extra 3% explained for the transient C1, 6% for the multifocal C1, 16% for the 18.75 Hz SSVEP and 6% for the 7.5 Hz SSVEP, compared with the V1-only model. In the three-area model, the beta coefficient for V1 was largest in all cases, being more than twice the magnitude of that for V2/V3 for all four VEPs (Table 1). The lower BIC values for the three-area model in each case indicated that these improvements in model fit justified the increased complexity.

**Figure 6:**
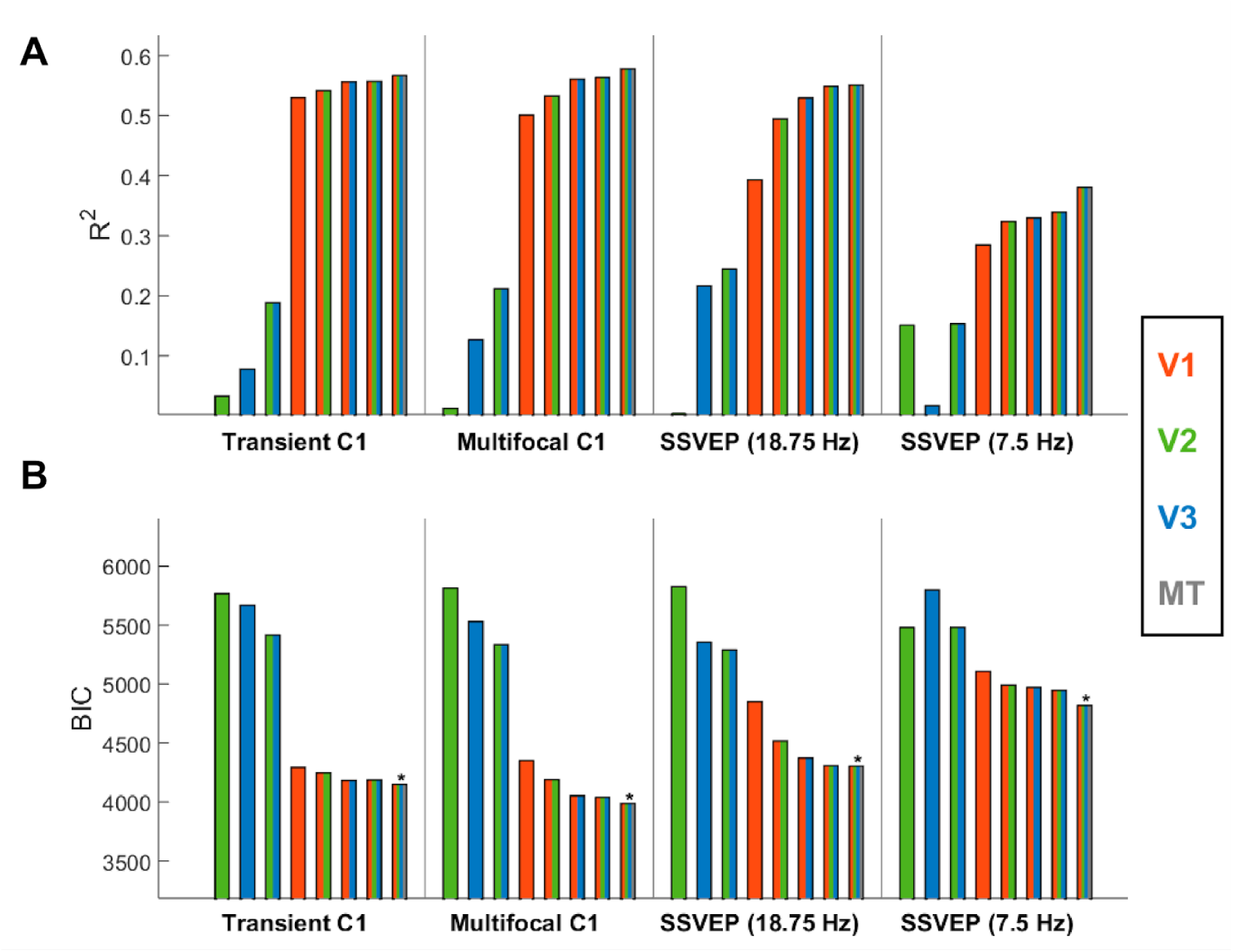
Regression models fitting each empirical VEP signal based on predicted topographies for V1, V2, V3 and MT, showing R^2^ (A) and BIC values (B). Each panel corresponds to a signal and each bar to a model, with the set of colours within each bar denoting the visual areas included. The model with the lowest BIC for each signal is marked with an asterisk.

**Table 1:**
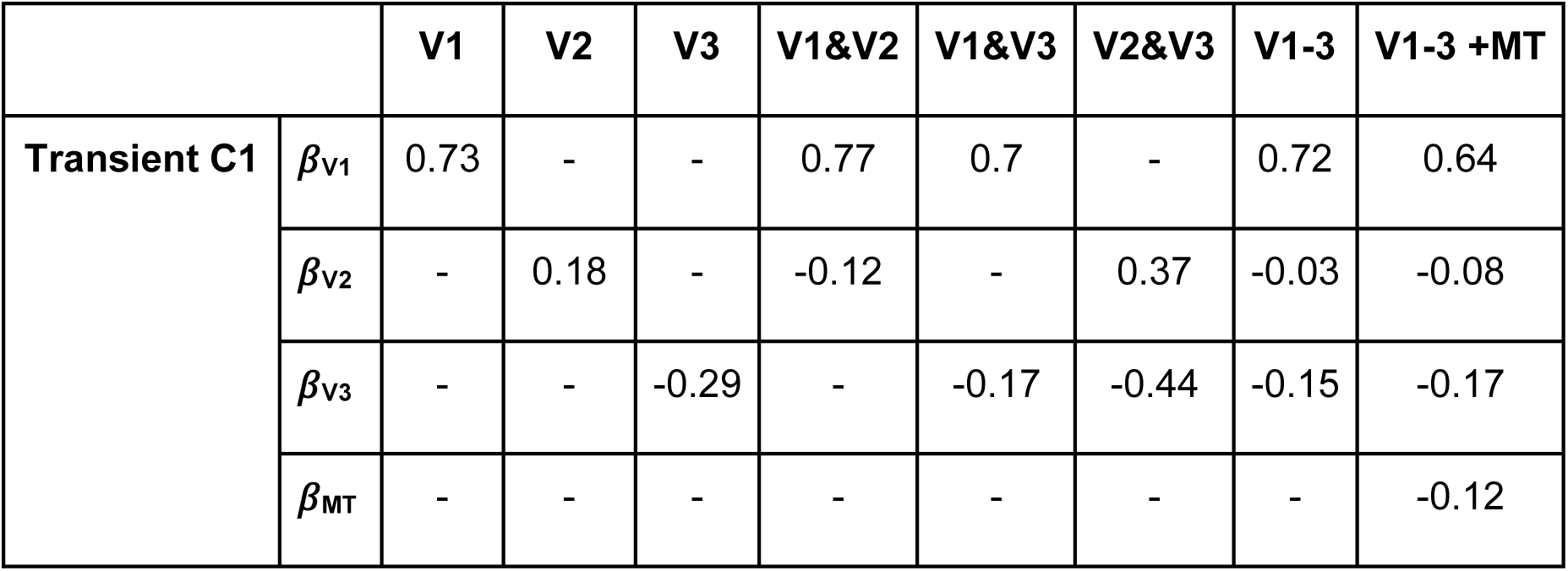
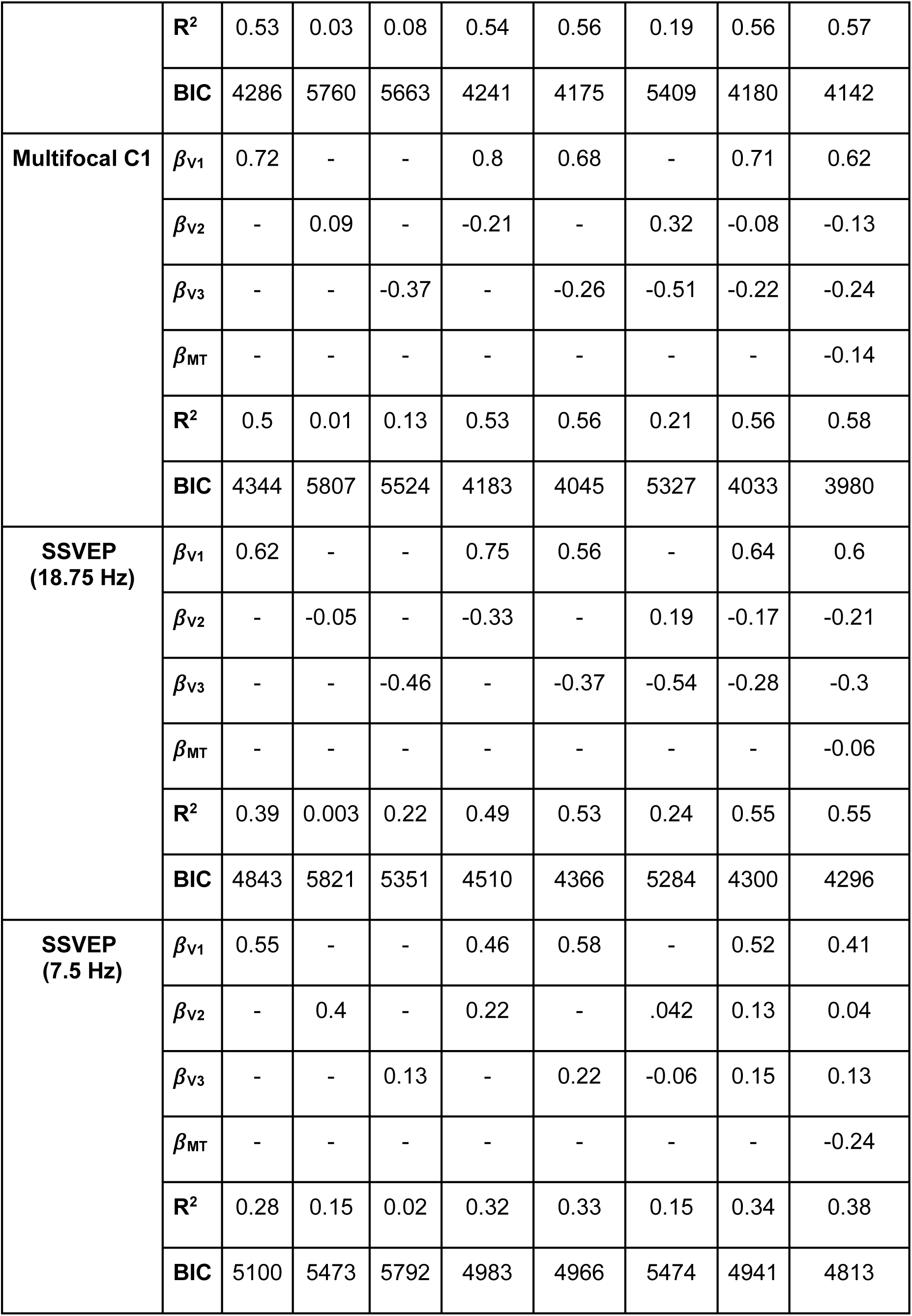
Beta coefficients, R^2^, and BIC values of linear regression models predicting empirical VEP signals from MRI-predicted topographies for V1-3 and MT.

So far, we have only considered visual areas V1, V2 and V3 because these are the areas implicated in the debate surrounding C1 generation (Ales et al., 2013; Kelly, Schroeder, et al., 2013; Kelly, Vanegas, et al., 2013) and because these are the only areas in the Benson-2014 retinotopy atlas known to reverse in polarity across the horizontal meridian. However, in one previous SSVEP source analysis study, a prominent contribution from MT was identified in addition to V1 (Di Russo et al., 2007). Therefore, to follow up, we fit a model including area MT in addition to V1, V2 and V3, though the results of it should be treated with caution because MT retinotopy carries considerably more uncertainty than V1-3 since the MT complex comprises a number of smaller sub-areas with larger receptive field sizes (Amano et al., 2009; Kolster et al., 2010; Pitzalis et al., 2010). This model explained 57% of variance in the transient C1 (increase of 1% compared with the V1-3 model) with an MT weight of −0.12, 58% (increase of 2%) in the multifocal C1 with an MT weight of −0.14, 55% (no change) in the 18.75 Hz SSVEP with an MT weight of −0.06, and 38% (increase of 4%) in the 7.5 Hz SSVEP with an MT weight of −0.24. This led to a drop in BIC compared to the V1-3 model in all cases, but the drop was largest for the 7.5 Hz SSVEP (Figure 6B, Table 1). Meanwhile, models with MT alone explained 11% of variance in the transient C1, 9% in the multifocal C1, 2% in the 18.75 Hz SSVEP and 22% in the 7.5 Hz SSVEP.

### 3.3 Temporal dynamics of model fits for the transient and multifocal VEP

The models for the transient and multifocal VEPs have thus far been static in the sense that they have modelled a single measurement window of 70-80 ms following stimulus onset to correspond with the C1 peak latency. However, to demonstrate that the results generalise beyond this chosen time window, we applied the modelling procedure to a series of 10-ms VEP measurement windows in 5-ms steps from −100 to 400 ms. This produces time courses of activation for V1, V2, V3 and MT, analogous to those derived previously using RCSE (J. Ales et al., 2010; Hagler, 2014; Hagler et al., 2009; Hagler Jr. & Dale, 2013) and they help to inform judgements about what time window to use to measure activity in a particular visual area. For example, it has been suggested that the rising slope of the C1 may be most appropriate to measure V1 activity without overlap from other visual areas (Foxe & Simpson, 2002; Plomp et al., 2010; Vanni et al., 2004). Tracing the relative dominance of the areas over time will provide further insight into this.

Figure 7 shows the time courses of R^2^ values and *β*-coefficients for both the single-area and four-area models of the transient and multifocal VEPs. Note that the scale of the *β*-coefficients is different to the earlier models simply because the signals were z-scored based on means and standard deviations from different measurement windows (70 to 80 ms before and −100 to 400 ms here). The time courses of R^2^ values indicated that the transient VEP began with a peak at 70-80 ms, consistent with the C1 component, that was mostly explained by V1 (Fig. 7A). This was also true of the mfVEP (Fig. 7D), although there were additional pre-stimulus R^2^ peaks, reflecting a small amount of auto-correlation in the multifocal pulse streams (*r=0.1*) incurred by the use of one out of every four pulses (see Methods). The initial peaks at 70-80 ms were followed by a sequence of recurring peaks, variably explained by striate and extrastriate areas (Fig. 7A&D). Correspondingly, the time courses of *β*-coefficients for both the single-area models (Fig. 7B&E) and full models (Fig. 7C&F) exhibited a sequence of positive and negative peaks in each visual area, the latencies of which are provided in Supplementary Tables S1-4. Although the dominance of V1 can be observed to persist through the C1 timeframe across the board, the nature and extent of contributions from areas V2, V3 and MT differ in the full models compared with the single-area models (compare Fig. 7B&E with Fig. 7C&F). In the single-area models, in which any variance possible must be attributed to that one area, the V1 model’s initial positive peak coincided with negative peaks in the V3 and MT models of almost half the magnitude and a delayed positive peak in the V2 model (Fig. 7B&E). In the full model, where variance is free to be attributed to any area to achieve best fit, V2 showed an initial negative peak alongside V3 and MT, and these were approximately one fifth the size of the concurrent V1 peak (Fig. 7C&F). By contrast, the size of the V1 peak did not undergo any major change between the single-area and full models. Inferring the true degree of extrastriate contributions is complicated by the fact that the predicted topographies for the different visual areas are correlated to varying extents (Table 2), which can allow one area to account for explained variance that is really due to another area, and this “cross-talk” can in principle occur in both the single-area and full models. However, here, a comparison across signal types can provide valuable insight. In the mfVEP, V1 remains the dominant contributor throughout the waveform (Fig 7D) and there exists at least one moment (e.g. approximately 120 ms) where all model coefficients pass through zero because there is no variance to pick up on (Fig 7D-F; Fig 9B). These features are highly inconsistent with a cascade of activity across multiple visual areas, and much more consistent with a multiphasic response emanating from a sole generator in V1 (Lalor et al., 2012; Slotnick et al., 1999). It would follow that any periods of non-zero amplitude in the extrastriate waveforms estimated from the mfVEP purely result from cross-talk due to the partial topographic correlations (Table 2). Consistent with this account, the non-zero extrastriate contributions during the C1 window (70-90 ms) in the single-area models of the mfVEP have the same sign as these correlations among predicted topographies, positive for V2 and negative for V3 and MT (Table 2; Fig 7E). Together, this all suggests that the extrastriate contributions in the mfVEP can serve as a benchmark for the degree of cross-talk that stands to occur in the transient VEP (Fig 7C). Notably, the extrastriate areas make similar initial contributions to the transient VEP as to the mfVEP, suggesting that this too is the product of cross-talk (Fig 7B&E). In both VEPs, these initial extrastriate contributions are also diminished in the multi-area model while V1 coefficients remain just as strong. Taken together, these comparisons suggest that cross-talk, whereby extrastriate areas account for V1-driven variations, occurs strongly in the single-area models where V1 is not itself included to account for them, and more weakly in the full models. Moreover, the estimated extrastriate contribution in the transient VEP appears commensurate with that expected from cross-talk according to the mfVEP during the C1 timeframe. However, in the transient VEP unlike the mfVEP, extrastriate contributions become clearly dominant beyond the C1 timeframe (Fig 7A-C).

**Figure 7:**
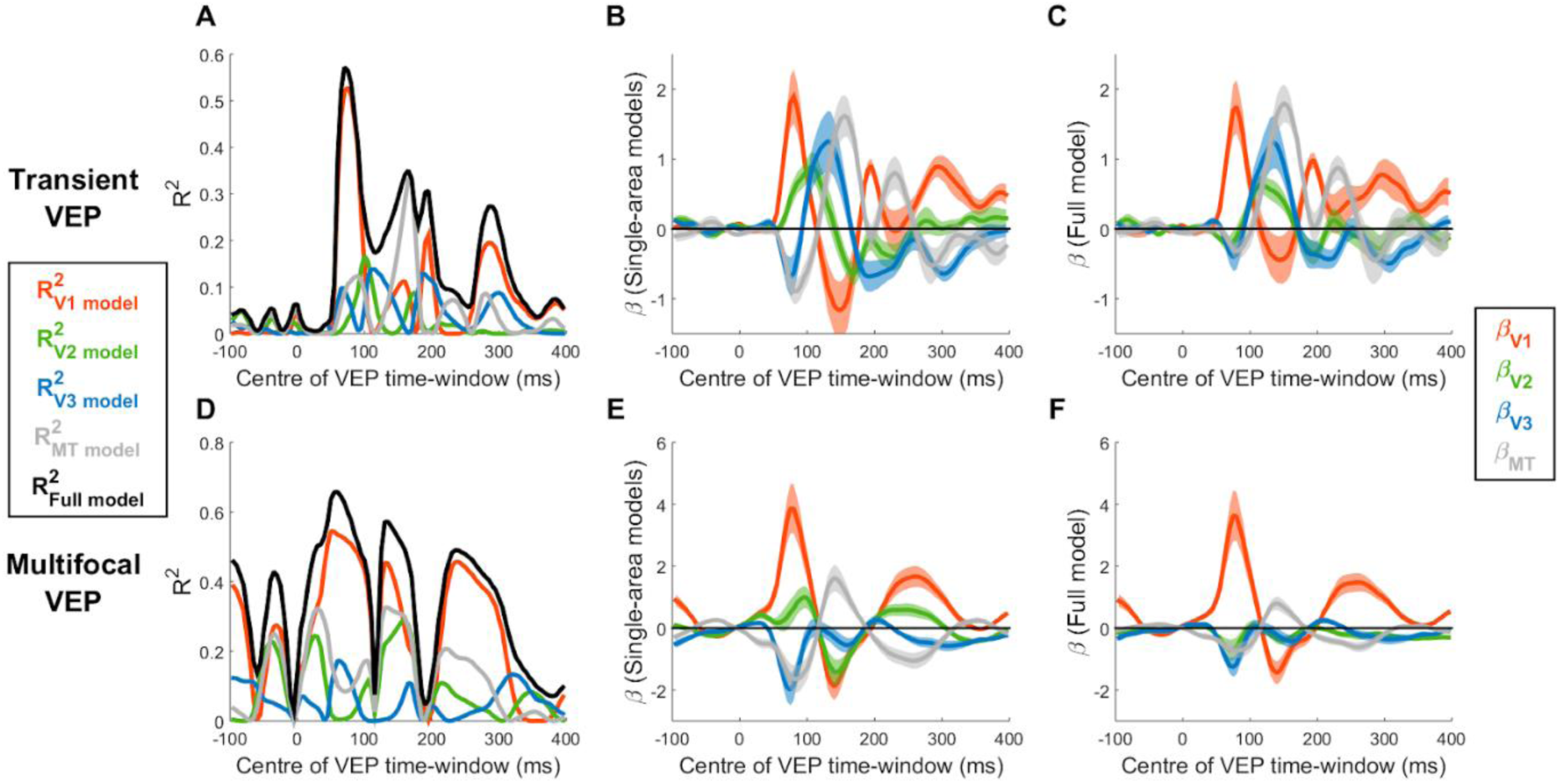
Model fits of the transient VEP (A-C) and the multifocal VEP (D-F) as a function of the centre-point of the 10-ms signal measurement window. A&D) R^2^ values across time for single-area models (V1, V2, V3 and MT) and the full model including all four visual areas. B&E) *β*-coefficients ±1 bootstrap standard error for each of these four visual areas in the single-area models. C&F) *β*-coefficients ±1 standard error for each of these four visual areas in the full model.

**Table 2:**
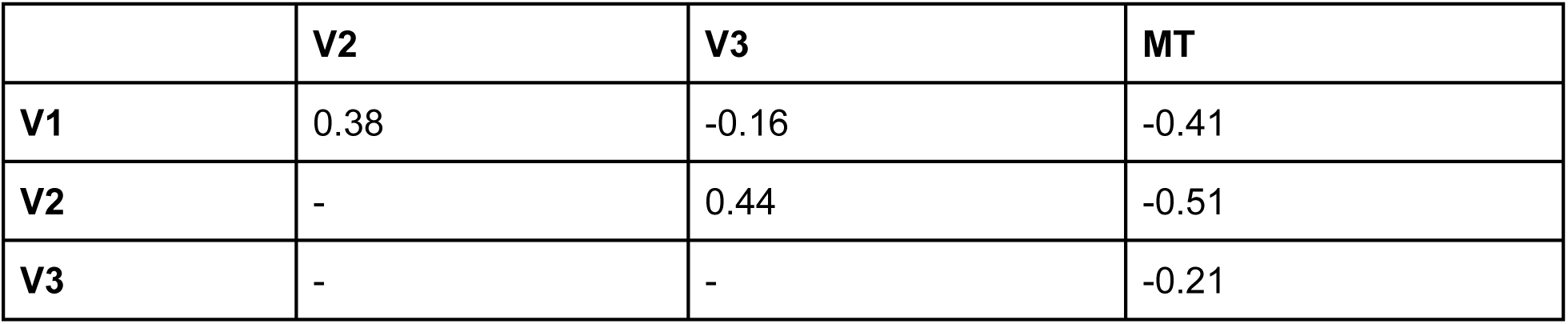
Pearson’s correlation coefficients among the MRI-predicted topographies of V1, V2, V3 and MT.

|  | V2 | V3 | MT |
| --- | --- | --- | --- |
| V1 | 0.38 | -0.16 | -0.41 |
| V2 | - | 0.44 | -0.51 |
| V3 | - | - | -0.21 |

In light of the dominance of V1 over V2, V3 and MT during the C1 time-frame, both in terms of variance explained and the magnitude of *β*-coefficients, we calculated the difference in absolute *β*-coefficients between V1 and each other area to determine the C1-measurement window where V1 contributions are maximally stronger. These are plotted along with V1 *β*-coefficients in Figure 8 for both the single-area models and full models, and both the transient and multifocal VEP. These plots suggest that V1’s superiority over other areas hits its peak at approximately the same time as V1 itself hits its peak, at approximately 70-90 ms in line with C1 peak latency. Assuming that the bulk of EEG noise does not scale with amplitude, this suggests that strategies of measuring V1 activity during the rising slope of the C1 to avoid overlap with other signals (Foxe & Simpson, 2002; Plomp et al., 2010; Vanni et al., 2004) may not achieve the best possible SNR of V1 responses.

**Figure 8:**
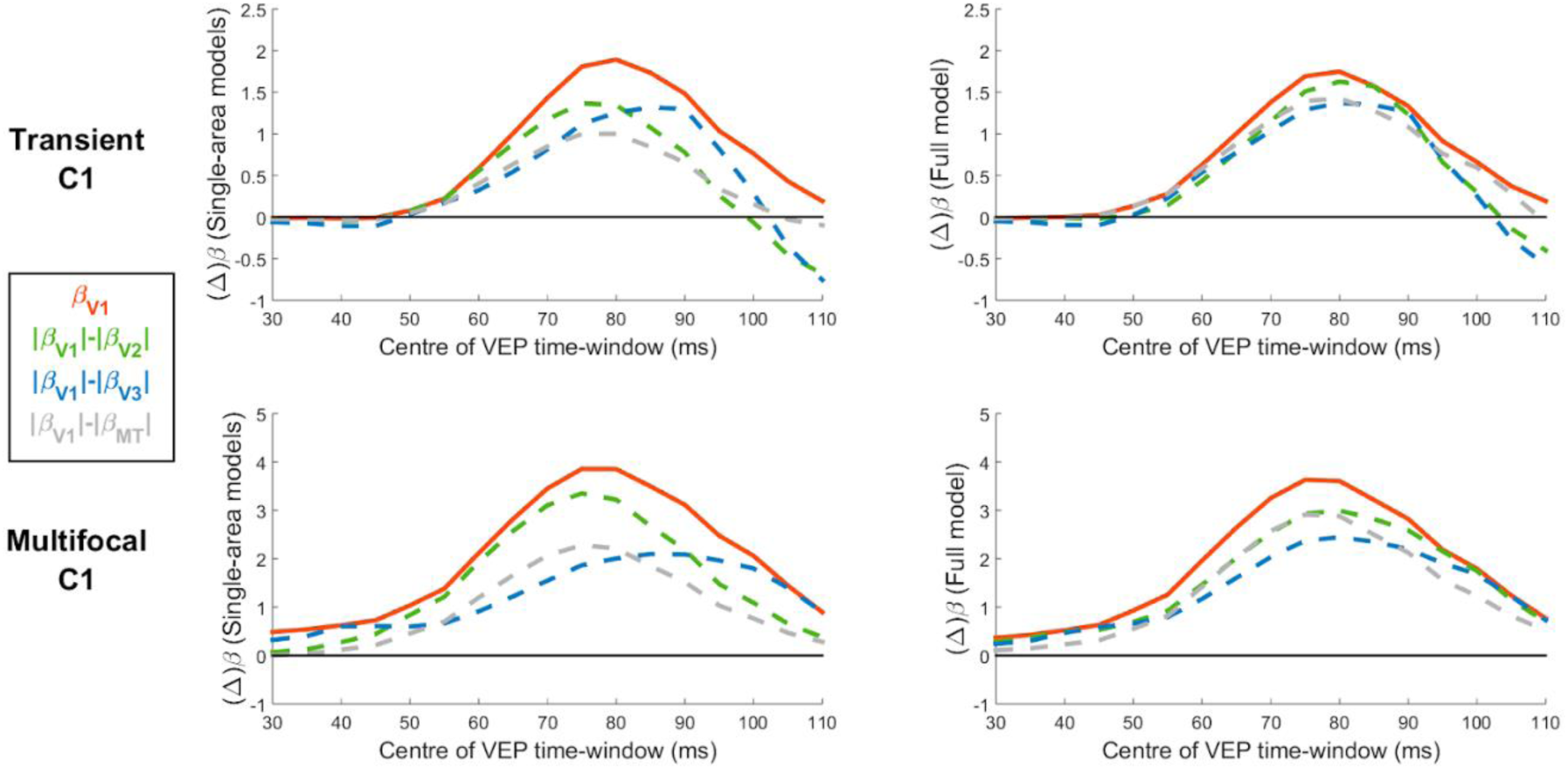
Difference in absolute *β*-coefficients of V1 relative to V2, V3 and MT during the C1 time frame for the transient and multifocal C1 and for single-area and full models. These *β*-differences are plotted alongside the *β*-coefficients of V1 itself to aid in the comparison of peak times, demonstrating that V1 *β*-coefficients are maximally different from those of the other three areas around the time of the C1 peak.

### 3.4 Phase shift in the 7.5Hz SSVEP

The only signal that did not qualitatively match the pattern of topographies predicted by V1 across the full visual field was the 7.5 Hz SSVEP, which deviated from the V1 pattern in the lower visual field (Figure 2D). This break in pattern was accompanied by a near-orthogonal shift in its phase between upper and lower visual field locations (see peak latency variability in the time domain, Figure 9D, and rotations in the complex plane, Figure 9E). This stood in contrast to the relatively stable phase of the 18.75 Hz SSVEP (Figure 9C&E) and a lack of clear variation in C1 latency across locations for the transient C1 (Figure 9A) and the multifocal C1 (Figure 9B). Topographies for the 7.5 Hz SSVEP at this orthogonal phase (calculated by projecting electrodes onto the plane orthogonal to the principal phase plane in Fig 9E) appeared more aligned to V2 or V3 (compare the inner circle of Figure 9F with Figure 2E-G). Thus, although the static models establish that V1 is the dominant generator of this slower SSVEP, a further analysis accounting for temporal dynamics in the SSVEP was warranted to explore the nature of secondary contributions from V2/V3.

**Figure 9:**
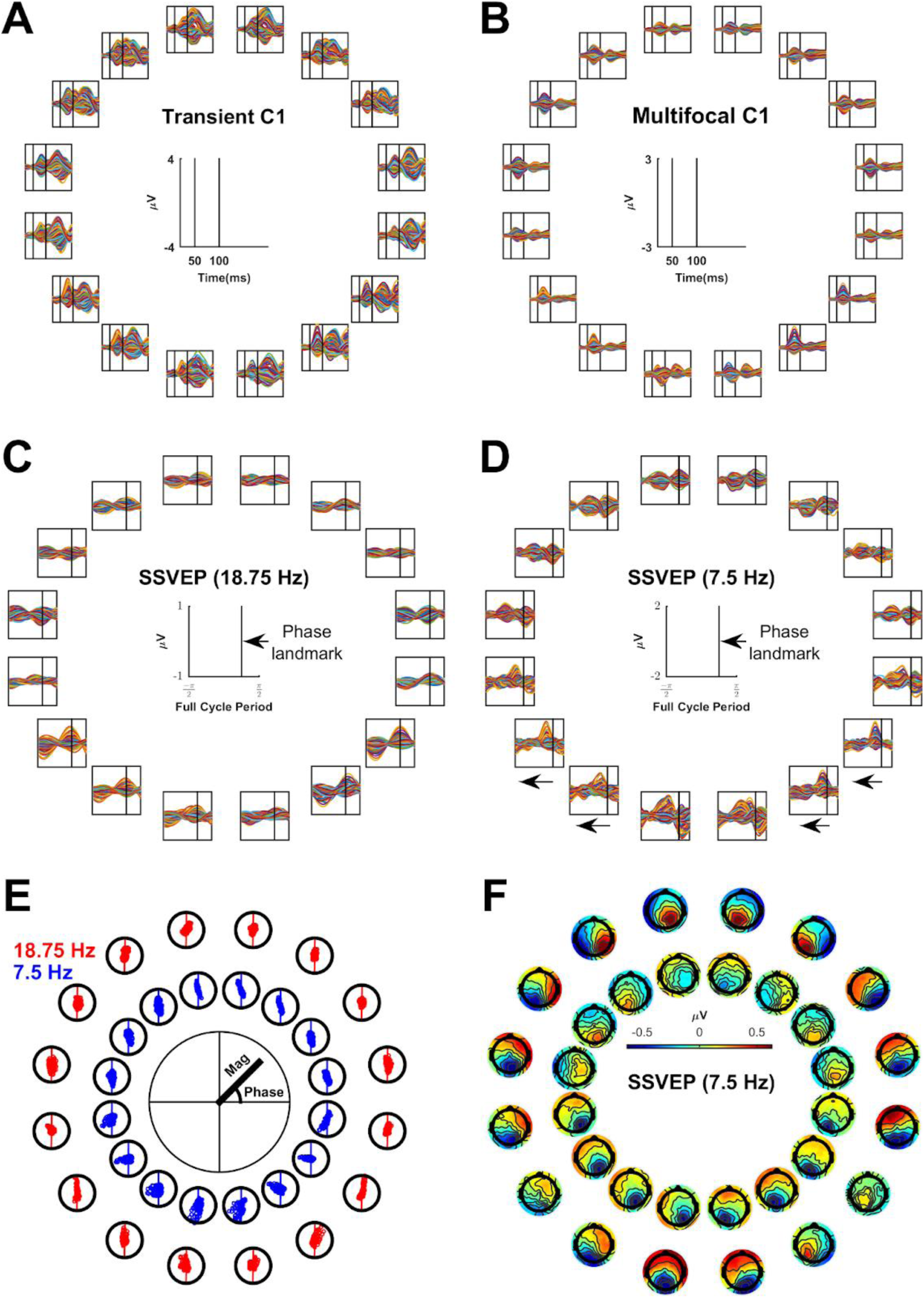
**A-B:** Butterfly plots of the transient and multifocal C1. Vertical black lines indicate latencies of 50 ms and 100 ms respectively, serving as temporal landmarks to aid comparison. **C-D:** Butterfly plots of a single cycle of the 18.75 Hz SSVEP and the 7.5 Hz SSVEP with black vertical lines acting as a temporal landmark to visualise the phase shift in the 7.5 Hz SSVEP. **E:** Scatter plots of complex FFT values for the 18.75 Hz and 7.5 Hz SSVEPs (each point corresponds to an electrode). Oscillation magnitude is given by the distance of each point from the centre while phase is given by its polar angle. Plots have been rotated so that the principal phase, i.e. the line best traversing the points, is aligned to the vertical axis. **F:** Topographies for the 7.5 Hz SSVEP are shown (outer ring) with the additional inclusion of topographies corresponding to a 90° clockwise phase shift (delay), calculated by projecting electrodes onto the axis orthogonal to the principal phase axis (inner ring).

To explore how this phase shift in the scalp-recorded signal emerged, we considered how oscillations originating in V1 and V2/V3 would sum together on the scalp. V1 and V2/V3 likely produce signals of opposite polarity on the scalp due to their opposing geometry. Without any transmission delays among them at the cortical level, this would simply lead to destructive interference with the larger signal prevailing. However, with a transmission delay at the cortical level we would additionally expect a change in phase at the scalp level because a classic trigonometric identity tells us that when two time-lagged oscillations of the same frequency are summed, the resultant phase is at the midpoint between the two original phases: cos[⍵(t)+п]+cos[⍵(t)-d]=2cos[(п+d)/2]cos[⍵(t)+(п-d)/2]. If the phase delay at the cortical level, d, is small, then the resulting phase shift at the scalp level is indeed close to 90° because (п±d)/2 is close to 90°. We reasoned that this dynamic could be at the heart of the phase shift because responses in V2/V3 likely do lag those in V1 (Givre et al., 1994; Mitzdorf, 1987; Schroeder et al., 1991, 1998; Xing et al., 2009). However, the above trigonometric identity assumes the summed oscillations have the same magnitude, whereas in reality, the signals in V1 versus V2/V3 likely differ in amplitude depending on visual field location (e.g. Ming et al., 2020), and even simply due to varying scalp proximity; for example, upper field stimuli project to ventral sites further from the scalp (Fig 1B-C; Fig 10A). What’s more, the empirical phase shift was observed in a specific case only: the lower visual field for the 7.5 Hz SSVEP, not the upper visual field or the 18.75 Hz SSVEP. Thus, the phase shift may depend not only on latency differences among V1, V2 and V3, but also on magnitude differences and oscillation frequency.

**Figure 10:**
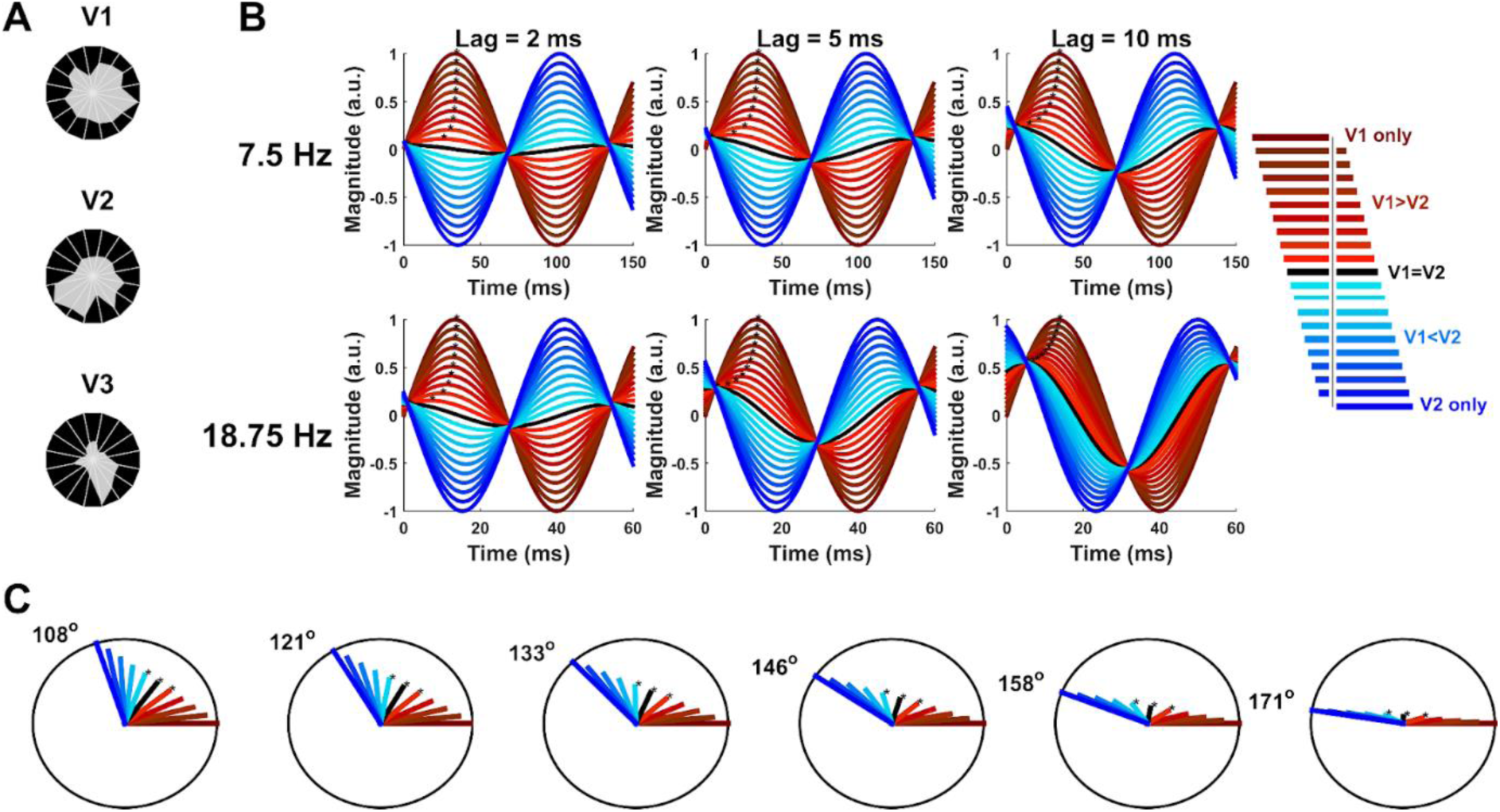
Simulations of sinusoids with opposing polarity for insight into the 7.5 Hz SSVEP phase shift effect. **A)** Magnitude of the MRI-predicted topographies (peak amplitude) as a function of visual-field polar angle, showing a greater difference in scalp projection strength between the upper and lower visual field for V2/V3 than for V1. Distance of the grey shading from the circle-centre represents magnitude at that polar angle. **B)** Sums of opposite-phase sinusoids representing oppositely-oriented V1 and V2 signals as a function of their oscillation frequency (top/bottom), relative weight (traces within panels) and latency delay (left to right). Red and blue curves correspond to various relative weights of V1 and V2; black curves correspond to equal weighting. The blue curve is delayed relative to the red curve and in the cases shown here, this results in a near 90° shift of the black curve earlier in time. Black asterisks outline the trajectory of peak latency for each signal weighting, demonstrating that the abruptness of the phase shift (as a function of change of weighting) depends on the frequency and latency delay. **C)** Sums of sinusoids shown on the complex plane. Note that whereas the time plots in B illustrated different latency delays and oscillation frequencies separately, these phasor plots combine them into phase delays. Also note that since an oscillation of opposite polarity equates to a phase difference of 180°, phase delays close to 180° in these phasor plots correspond to small latency delays in panel B while phase delays further from 180° correspond to larger latency delays. Different magnitudes of V1/V2 are represented by the same colour scheme as panel B. Asterisks highlight the equal magnitude case and the two flanking cases in which either the V1 or V2 oscillation is slightly larger. Six different phase delays are shown, demonstrating that the phase of the sum shifts as a function of relative magnitude, and does so more abruptly near the case of equal-magnitude for phase delays closer to 180° (i.e. those with shorter time delays between oscillations of opposite polarity).

In the section to follow we present a regression analysis to estimate the time lags and relative magnitudes among the visual areas that could explain the magnitude and phase of both oscillation frequencies on the scalp, with the goal of explaining why the large phase shift occurs selectively in the 7.5 Hz SSVEP. To first provide an intuition for how time lags, relative magnitudes and oscillatory frequency interact to determine phase-shifting behaviour, we simulated sums of sinusoids, shown in both time-domain plots (Fig 10B) and phasor plots (Fig 10C). These simulations demonstrate that the degree to which a sum of sinusoids shifts in phase is a function of their relative magnitude and phase delay, where phase delay is jointly determined by time lag and oscillation frequency. At the extremes of complete dominance of V1 or V2, the phase of the sum is entirely aligned to that of the dominant area, and as the relative magnitude shifts from one area to the other, the phase of the sum shifts monotonically from one to the other extreme. At phase delays close to 180° (right-most plot in Fig 10C), the bulk of this phase shift takes place within a tight range of relative magnitudes close to equality, with little additional change in phase as one oscillation becomes much larger than the other. By contrast, at phase delays further from 180° (the left-most panel in Fig 10C), phase shifts more gradually across a broader range of relative magnitudes. Equivalently, in Fig 10B it can be seen that for a given time delay, phase shifts more gradually across a broader range of relative magnitudes for the 18.75 Hz oscillation than for the 7.5 Hz oscillation. This is because for faster oscillations, a given time lag is a larger phase delay.

These dynamics offer a potential explanation for the pattern of phase shifts observed in the SSVEP. That it took place between the upper and lower visual field could be explained by small changes in relative magnitude between V1 and V2/V3 across this region because the simulations demonstrated that small magnitude changes can in some cases produce large phase shifts. Moreover, that it was seen in the 7.5 Hz SSVEP and not the 18.75 Hz SSVEP could be the result of a smaller phase delay between V1 and V2/V3 for the 7.5 Hz SSVEP, leading to a larger shift in phase between the upper and lower visual field.

### 3.5 Modelling the 7.5 Hz phase shift

The simulations above suggested that the phase shift in the 7.5 Hz SSVEP between the upper and lower visual field could, in principle, be explained by time-lagged oscillations among V1, V2 and V3. To test this idea, we extended the earlier regression models with a sinusoidal component to capture SSVEP phase. In this model, V1, V2 and V3 followed sinusoidal time courses, each with its own phase, allowing them to lag one another. In order to investigate whether the same time lags among visual areas could produce the observed phase variations in the 7.5 Hz SSVEP simultaneously with the lack of phase variation in the 18.75 Hz SSVEP, the two SSVEPs were modelled in tandem with shared time lags. We did, however, allow magnitude to vary between the SSVEP frequencies to allow for the possibility that response magnitude from visual areas may vary as a function of stimulation rate. Since SSVEPs are not strictly sinusoidal (Norcia et al., 2015), we used the Fourier transform estimates of the SSVEPs to transform them into their equivalent sinusoidal form for the purpose of model fitting. This is justified since the purpose of this modelling effort was not to explain deviations in the SSVEPs from sinusoidal oscillation and so this simply serves to remove a source of noise that is not centrally relevant to the modelling goal.

The structure of the data for these regressions is outlined in Figure 11. As before, all regressors and dependent variables were converted to z-scores before carrying out the regressions. To construct the dependent variable, a single cycle of a sinusoidal time-series was generated from the observed Fourier magnitude and phase at the appropriate SSVEP frequency for each electrode, retinotopic location and SSVEP frequency, and these time-courses were stacked in a single vector to model them together (see Fig 11A-B). A predictor variable was then generated in the same way for each visual area and both SSVEPs (6 total), with values for a given predictor set to 0 for observations related to the other SSVEP (Fig 11B). This regression analysis was repeated for a series of time lags for V2 and V3 relative to V1 in steps equal to the EEG sample rate (approx. 2 ms) in the range ±66 ms (half the cycle duration of the 7.5 Hz SSVEP). Because the “baseline phase” of V1 itself is unknown a priori and has no predictable relationship across different oscillatory frequencies, we also manipulated V1’s baseline phase independently for the two SSVEP frequencies across separate regression models, spanning a full cycle in both cases. Thus, while each regression model had 6 beta coefficients to capture relative amplitudes, the full range of combinations of the two baseline V1 phases, the V2 time lag and the V3 time lag (both relative to V1) were probed across separate regressions. This formulation constrained time lags to be the same for the two SSVEP frequencies while also allowing their baseline phases to differ. The regression model with the highest R^2^ out of these four manipulated phase/time dimensions was identified, and the estimates of all 10 relevant parameters taken from that model.

**Figure 11:**
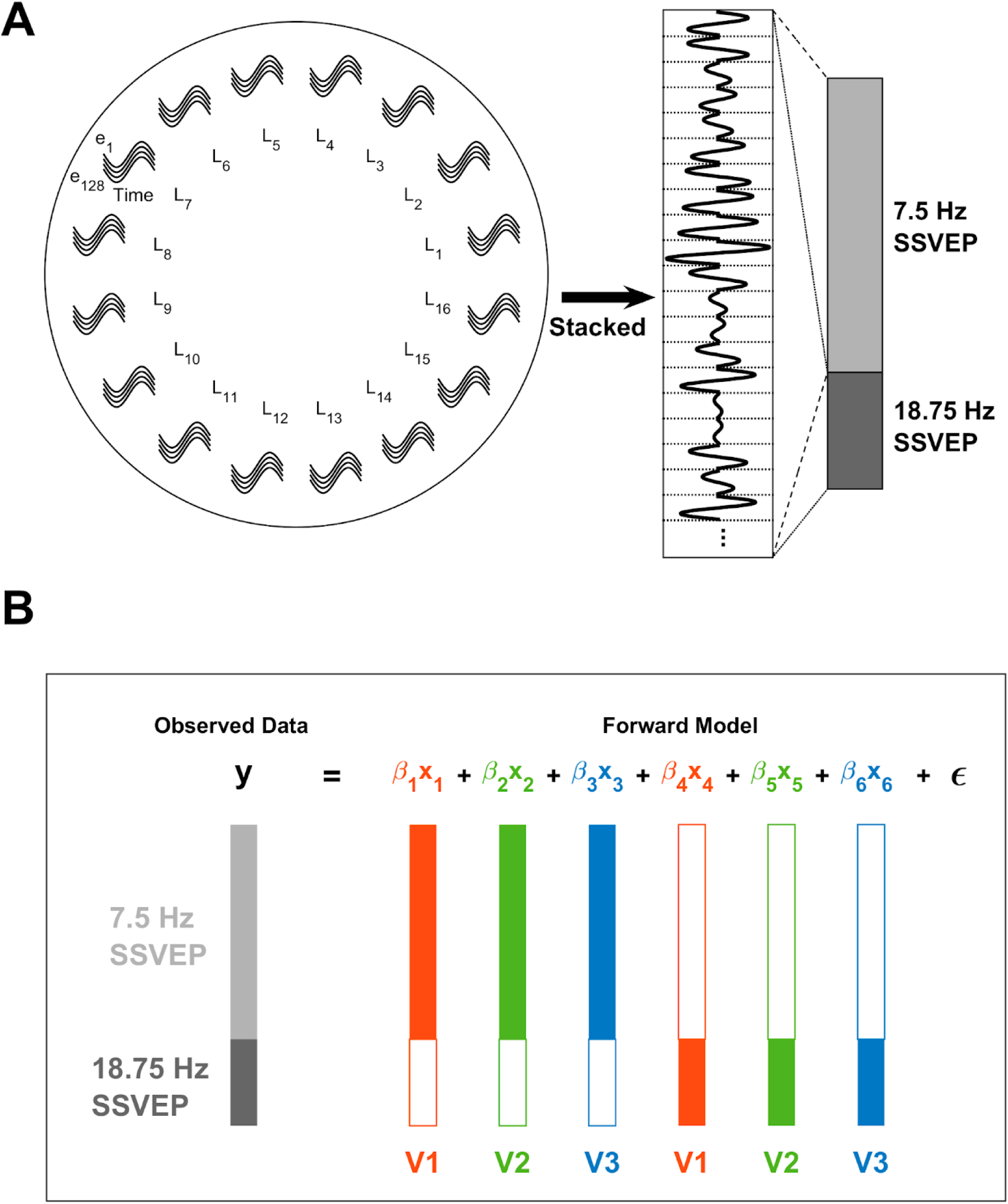
Diagram illustrating the structure of the sinusoidal regression models. (A) The structure of the observed data and the MRI-predicted regressors entered into the regression follows a common pattern: For each frequency, the phase and magnitude of the observed or predicted SSVEP was used to generate a single-cycle, sinusoidal time series for each electrode (e) and visual field location (L_n_). These sinusoidal snippets were then stacked for both the slow and fast SSVEP frequencies. To form the dependent variable, the phase and magnitude of the sinusoidal snippets were taken from Fourier analysis of the observed EEG. To form visual area regressors, the sinusoidal snippets were generated by using the MRI-predicted magnitudes at each electrode (e) and visual field location (L_n_) from the normalised forward model topographies (Fig. 2E-G) to scale single-cycle sine waves that are phase-shifted in accordance with the assumed baseline phases and time lags in each regression model (see main text). (B) In the regression model, the observed data vectors for the two SSVEP frequencies were further stacked to model them together. For each regressor, vectors for the two SSVEP frequencies were also stacked to align with the dependent variable even though each regressor pertained to only one of the two SSVEP frequencies, with values corresponding to the other SSVEP frequency set to zero. Note that the different lengths of bars correspond to the different number of data points in each SSVEP due to the longer cycle lengths of lower frequency oscillations.

Because baseline phases, V2/3 time lags and V1-3 magnitudes all varied independently, models could emerge with different combinations of these variables that were actually equivalent, as reflected in the identity cos[A]=-cos[A+*π*]. For ease of interpretation therefore, only models with all-positive magnitudes were considered. The distribution of R^2^ values, time lags and magnitudes among the top 1% best-fitting of these models is shown in Figure 12. The overall best-fitting model is shown by larger circles within these distributions. This model had a lag of 20 ms for V2 and 28 ms for V3. Its weights were highest for V1 for both the 18.75 Hz SSVEP (V1: *ꞵ=0.83*, V2: *ꞵ=0.36*, V3: *ꞵ=0.35*) and for the 7.5 Hz SSVEP (V1: *ꞵ=0.70*, V2: *ꞵ=0.17*, V3: *ꞵ=0.42*). Overall, this model had an R^2^ of 0.41, and breaking this down by SSVEP gave an R^2^ of 0.47 for the 18.75 Hz SSVEP and an R^2^ of 0.39 for the 7.5 Hz SSVEP (note, the skew here is because 71.5% of the observations to be fit came from the 7.5 Hz SSVEP due to the different cycle lengths). By comparison, the best model out of those where V2 and V3 lags were 0 ms (analogous to the earlier static models) had an R^2^ of 0.27 for the 7.5 Hz SSVEP and 0.43 for the 18.75 Hz SSVEP, reflecting improvements in the overall best model of 0.11 and 0.04, respectively. As shown in Figure 13, this improvement in model fit, which was particularly high for the 7.5 Hz SSVEP, was achieved by capturing the qualitative phase dynamics of the empirical SSVEPs. The model generated phase shifts in the lower visual field for the 7.5 Hz SSVEP (Fig. 13B&D), culminating in a delayed signal (Fig. 13B), while no major phase shifts appeared in the 18.75 Hz SSVEP (Fig. 13D). There were some discrepancies however. In the empirical data, the bulk of the phase shifts took place in the 2nd and 3rd locations from the lower vertical meridian (Figure 13A) whereas in the model they took place in the 3rd and 4th locations (Figure 13B). Moreover, orthogonal to the principal SSVEP phase, the empirical data demonstrated a disparity in magnitudes between the upper and lower visual fields (Figure 13E), being approximately 3 times larger in the lower field, while this difference was less obvious in the model (Figure 13F). In fact, the difference in the model was comparable in magnitude to upper-lower field differences seen in the empirical 18.75 Hz SSVEP (approximately 2:1) and so may simply reflect general magnitude differences across hemifields rather than the exaggerated difference brought about by phase shifts in the empirical 7.5 Hz SSVEP.

**Figure 12:**
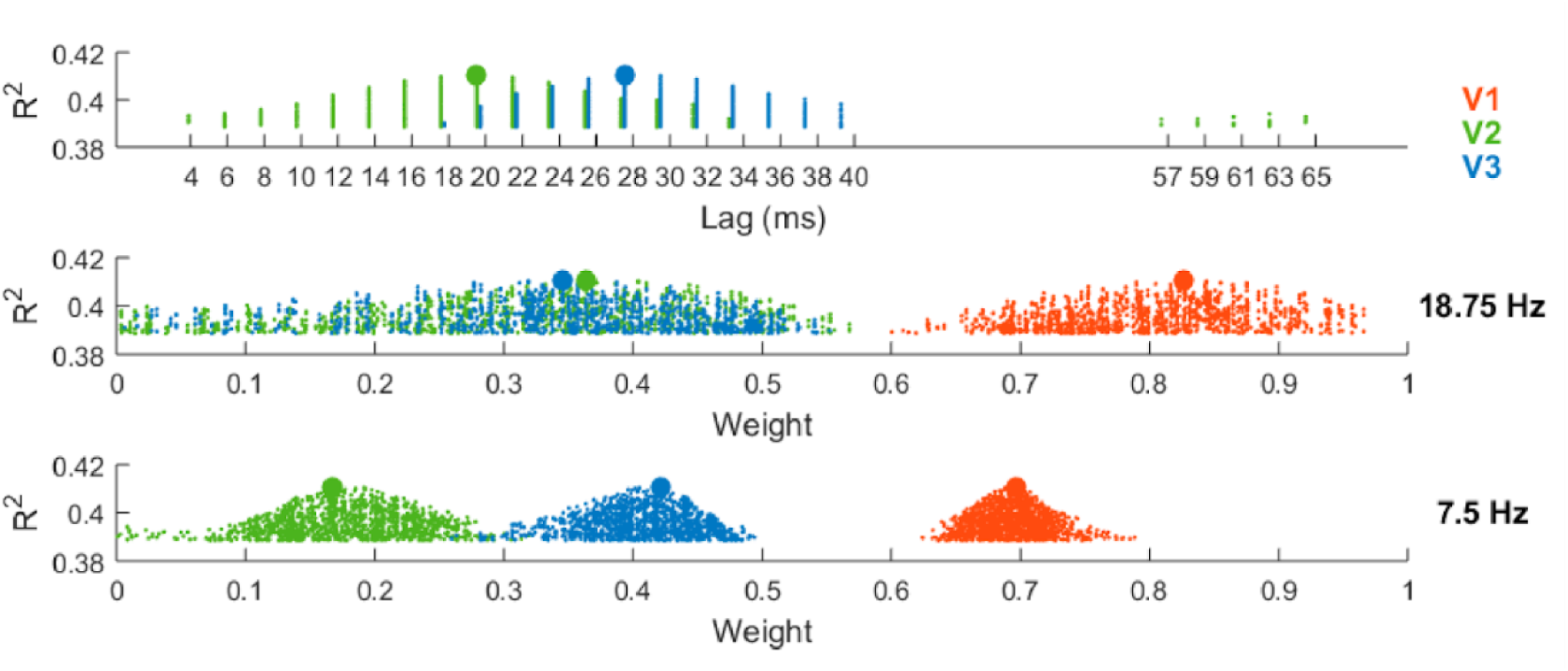
Explained variance of the regression models fit to sinusoidal time-courses of both SSVEP frequencies, aimed at explaining the observed 7.5 Hz SSVEP phase shift (Fig 2D). R^2^ values are plotted against the time lags (top) of V2 (green) and V3 (blue) relative to V1 for the top 1% best fitting regression models. Each dot represents one model and the two larger dots correspond to the overall best model. Below, similar parameter distributions are plotted for weight (V1-3) in the 18.75 Hz SSVEP and the 7.5 Hz SSVEP.

**Figure 13:**
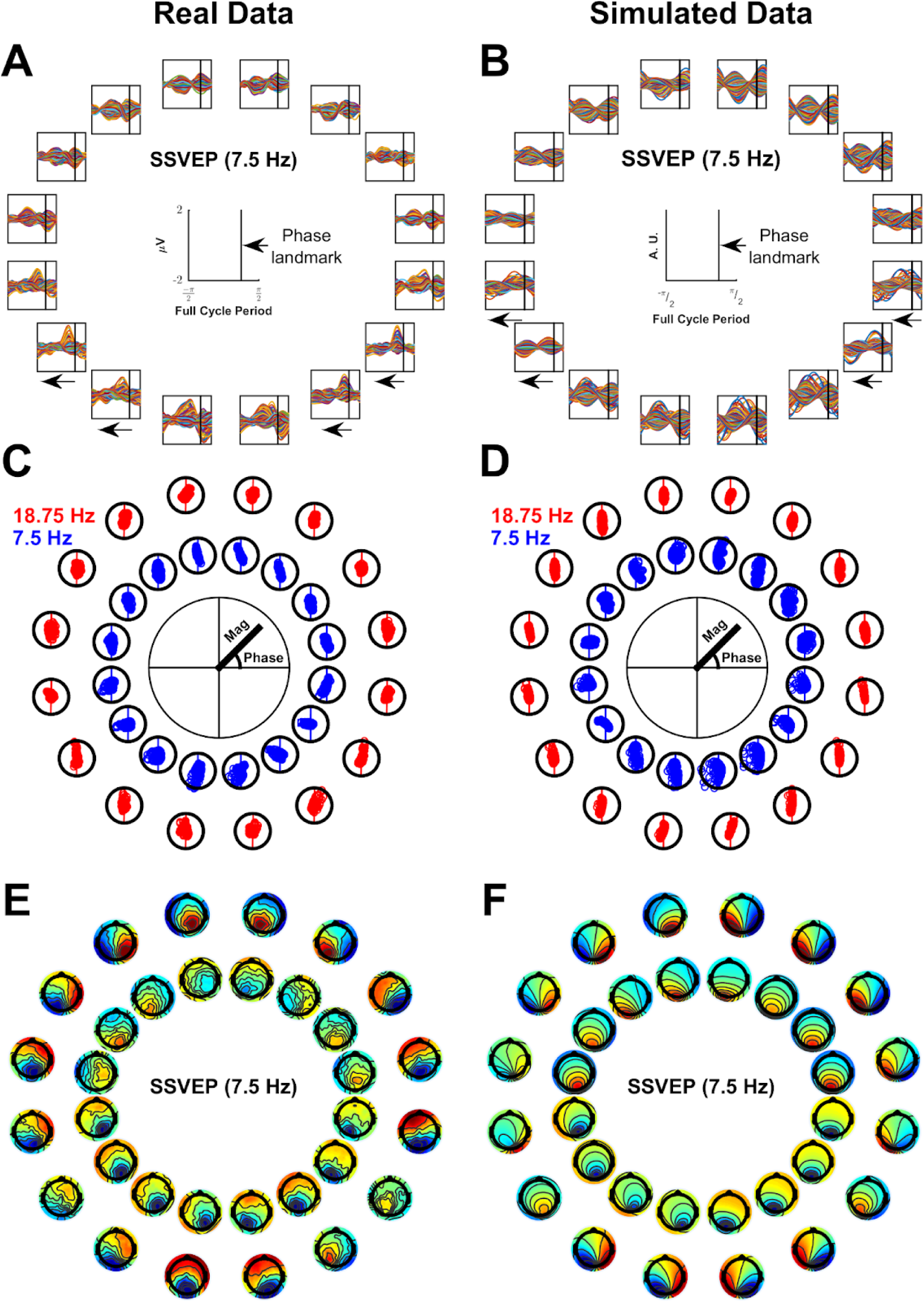
Real Data (A, C, E) and simulated data (B, D, F) for the overall best sinusoidal model. **A-B)** Butterfly plots of a single cycle of the 7.5 Hz SSVEP with black vertical lines acting as a temporal landmark to visualise phase shifts. **C-D)** Scatter plots of the 18.75 Hz and 7.5 Hz SSVEPs (each point corresponds to an electrode) demonstrating magnitude and phase. In the same way for the real and simulated data, the phase axis was rotated so that the principal phase axis was vertical (shown by vertical lines traversing the points). **E-F)** Topographies for the 7.5 Hz SSVEP (outer ring) are shown with the additional inclusion of topographies corresponding to a 90° clockwise phase shift (delay), calculated by projecting electrodes onto the orthogonal phase (inner ring).

As with the static model, a follow-up model was carried out with MT additionally included in light of its previous implication in the SSVEP (Di Russo et al., 2007). Baseline phases and time lags were taken from the original model and fixed, allowing only the time lag of MT and the magnitudes of all four visual areas to vary. This was done for reasons of computation time given the combinatorial nature in which time lags were assigned to visual areas across regression models. Overall R^2^ increased from 0.41 to 0.43. This increase was achieved primarily through an improved fit of the 7.5 Hz SSVEP, for which R^2^ increased from 0.39 to 0.41, while R^2^ for the 18.75 Hz SSVEP remained at 0.47. MT lagged V1 by 8 ms and its weight was −0.13 for the 18.75 Hz SSVEP (compared to 0.78 for V1) and −0.26 for the 7.5 Hz SSVEP (compared to 0.62 for V1).

## 4 Discussion

Visual evoked potentials (VEPs) are widely used in Cognitive Neuroscience to gain insight into a range of brain operations from low-level visual encoding to higher-order cognitive processes. These various operations are studied using a mix of three complementary VEP techniques, and it is vital to determine the shared and distinct cortical sources of these VEP signals if we are to fully understand those operations and how they relate to one another. The goal of this study was to shed new light on the cortical origins of three low-level visual signals, the C1 component of the transient and multifocal VEP and the steady-state VEP (SSVEP), elicited at a fast (18.75 Hz) and slow (7.5 Hz) flicker rate. Using regressions of grand-average data against MRI-predicted scalp topographies for stimuli across the visual field, we found that V1 provided a substantially better account of scalp data than V2 or V3 for all four signals, and outlined a set of key V1-predicted topographic features which underlie this result. While the explained variance of the VEP data was incrementally improved with the addition of each of a series of extrastriate regions, multi-area models indicated that the bulk of extrastriate activations come substantially later in time, with those contributions emerging distinctly after the C1 peak in the transient VEP, with a most probable time lag of 20-30 ms in the SSVEP, and potentially not emerging at all in the multifocal VEP. In the following we recap and integrate the findings across the VEP signals and analyses to lay out the basis for these conclusions.

Our forward-modelled topographies for V1, V2 and V3 predicted polarity inversion across the horizontal meridian in all cases, consistent with the forward models of J. M. Ales et al (2010), thus bolstering their conclusion that polarity inversion alone is an insufficient criterion for identifying a V1 source. However, above and beyond this polarity inversion, all four of the empirical VEP signals showed components of topographic variation both between and within visual field quadrants that each strongly distinguished V1 from extrastriate sources. Specifically, topographic foci were ipsilateral in the upper visual field and contralateral in the lower visual field, becoming more so as stimuli approach the vertical meridian. This is consistent with previous C1 reports (Clark et al., 1994; Di Russo et al., 2002) and also with simulations carried out by Ales et al (2013). Only V1 could account for the within-quadrant topography shifts because its extension from the Calcarine sulcus out onto the medial wall is uniquely consistent with them (Kelly, Vanegas, et al., 2013). V1 also captured the between-quadrant variations of the VEP signals better than did V2 or V3. The V1-predicted topographies were ipsilateral in upper visual field locations and contralateral in lower visual field locations, like the VEP signals, and this pattern of lateralization was reversed in V2 predictions while V3 predictions showed little change in lateralization across visual field quadrants. These broad lateralization trends across visual quadrants offer an important amendment to the polarity-inversion heuristic that is often used to diagnose a V1 source for VEP signals. Because V2 and V3 also predict this polarity inversion (J. M. Ales et al., 2010), the heuristic should further assert that signal polarity be slightly ipsilateral in the upper visual field and contralateral in the lower visual field to make it truly V1-specific. This simple heuristic offers a means to distinguish a V1 source from V2/V3 sources with some confidence based on stimuli in only one upper and one lower visual field location, as is commonly available in cognitive and perceptual studies. It should be noted, however, that this applies to the grand average and not to individual subjects, the topographies of whom are much more heterogeneous.

Overall, V1 explained approximately five times as much variance in the transient and multifocal C1 components and twice as much variance in the SSVEPs as V2/V3 and had regression weightings that were approximately four and three times stronger when visual areas were modelled together. Thus, among areas V1, V2 and V3, V1 clearly makes the dominant contribution to these VEP signals. However, V2, V3 and MT did explain significant additional variance when included alongside V1, raising the question of whether any of these signals are sufficiently isolated from extrastriate contributions to serve as a proxy for V1 activity measurements. This question has been of longstanding concern, particularly for the C1 component, and comes down to the relative strength and latencies of activation across visual areas. Although many animal electrophysiology studies find overlapping neural response latencies across visual areas, they tend to find an average temporal delay between responses in V1 and extrastriate areas (Bullier & Nowak, 1995; Chen et al., 2007; Maunsell & Gibson, 1992; Nowak et al., 1995; Raiguel et al., 1989; Robinson & Rugg, 1988; Schroeder et al., 1998). Previous attempts to model activation time courses of visual areas in humans using retinotopically constrained source estimation (RCSE) have yielded inconsistent results, with some finding simultaneous activation of V1 and V2 (J. Ales et al., 2010) and others finding that V2 and V3 both lag V1 (Hagler, 2014; Hagler Jr. & Dale, 2013). This discrepancy may be in part due to cross-talk that persists despite rich retinotopic constraints that are applied to each visual area. Our own time-resolved models of transient and multifocal VEPs found simultaneous initial activations across visual areas but there are a number of reasons to suspect that this is due to cross-talk. In fact, converging results across these and the SSVEP models are instead suggestive of extrastriate activations that peak at a considerable lag to V1. In support of the cross-talk account for the initial extrastriate activations, they were stronger in single-area models than multi-area models and were also congruent with the signs of correlations among predicted topographies of the visual areas. Moreover, key features of the multifocal VEP were strongly indicative of a sole source in V1 rather than a cascade of multiple areas, including the presence of timepoints following initial activation where all measures of activity passed through zero and the absence of recurrent V2 and V3 activations, suggesting that any estimated non-zero extrastriate contributions could be attributed to cross-talk. The transient VEP showed estimated extrastriate contributions that matched these inferred cross-talk contributions in polarity and magnitude during the C1 timeframe, whereas the later, first positive peaks of V2, V3 and MT were more consistent with true responses as they did not change between single- and multi-area models and were absent in the multifocal VEP. These positive-polarity peaks indicated response lags of 35 ms, 55 ms and 70 ms, respectively, relative to V1. Long V2 and V3 lags were also predicted by models of the SSVEP (20 ms and 28 ms), where they selectively accounted for a shift in phase that was specific to the lower visual field of the 7.5 Hz SSVEP and substantially improved on a zero-lag model. Although these lag estimates are not particularly close (35 vs 20 and 55 vs 28 ms), the shorter lags for the SSVEP are consistent with its greater improvement in model fit with inclusion of extrastriate areas and it is possible that the steady-state nature of the SSVEP facilitates shorter delays across visual areas. Overall, the level of agreement between the transient and steady-state models in pointing to a substantial extrastriate lag is remarkable because these were different signals evoked by different kinds of stimulation and the V2 and V3 lags explained an empirical phenomenon in the SSVEP that did not pertain to the transient or multifocal VEP (the phase shift). Thus, even allowing for some uncertainty in the precise estimation of these activation lags, our data from multiple stimulation protocols converge on extrastriate lags that are considerably far from 0 ms. The implication for the transient VEP is that there is a stretch of time on the order of tens of milliseconds during which V1 is the dominant contributor. Moreover, an analysis of the relative strength of beta weights over time indicated that the C1 peak represents the best measurement window for V1 activity in terms of signal-to-noise ratio.

While the above considerations suggest a considerable temporal delay between V1 and extrastriate visual areas in both the transient VEP and the SSVEP, V1 is not as easy to isolate in the SSVEP. Whereas in the transient VEP one can simply take the C1 time window to measure V1 activity, it is not clear that one can choose a particular phase of the SSVEP to achieve the same thing. The nature of time-lagged oscillations to combine into a single oscillation when summed may fundamentally limit our ability to separate visual sources in the SSVEP even if they oscillate at different phases as indicated in this study. Consistent with this, the “static” regression models suggested considerable contributions from V2/V3 in the SSVEP (between half and a third of the amplitude of V1 and adding between 6-16% explained variation). That extrastriate areas contribute to the SSVEP is also strongly implicated by our finding that time-lagged oscillations across visual areas could account for phase shifts between visual field locations and stimulation frequencies in the SSVEP. Di Russo et al (2007) also found in their source modelling work that the SSVEP was generated in both striate and extrastriate sources such as MT. One notable aspect of our results to compare with this is that we also found MT contributions in the 7.5 Hz SSVEP, a frequency almost as low as used in that study (6 Hz), but not in the 18.75 Hz SSVEP. This may reflect differing levels of MT involvement at different SSVEP frequencies but further investigation will be needed to address this question more comprehensively. Compared with the SSVEP, we found more modest extrastriate contributions to the C1 (explaining an additional 3% variance in the transient and 6% in the multifocal VEP). With a comparably greater extrastriate contribution to the SSVEP than the C1, comparisons between these two signals should always be made with this caveat in mind. One example in the literature that underscores this point is the divergence between the C1 and the SSVEP in terms of spatial attention. While SSVEP magnitude routinely increases with spatial attention (Morgan et al., 1996; Müller et al., 1998; Toffanin et al., 2009), similar effects in the C1 are much more rare (Hillyard & Anllo-Vento, 1998; Mohr et al., 2020; Qin et al., 2022). In addition to potential differences in recurrent activity contributions, the stronger extrastriate contribution to the SSVEP may partially explain this discrepancy, since higher level areas tend to show more robust spatial attention effects (Treue, 2001).

The modelling approach taken in this study was rooted in the cruciform model, determining cortical sources for visual signals based on the geometry of visual cortical areas. This approach shares similarities with a recently developed method - retinotopically constrained source estimation (RCSE) - that uses individual structural and/or functional retinotopic maps derived from MRI to constrain dipole models (J. Ales et al., 2010; Cottereau et al., 2015; Hagler, 2014; Hagler et al., 2009; Hagler Jr. & Dale, 2013; Inverso et al., 2016). This method was developed to address the issue in source analysis of non-unique source estimates and, to our knowledge, represents the state of the art of source analysis in the visual cortex. It does this by addressing the central reason why estimated sources are uncertain: neighbouring cortical areas that have similar aggregate surface orientation can masquerade as one another, making it impossible to distinguish them fully. By leveraging the assumption that visual responses in a visual area will be similar at different locations in the visual field, one can model a single visual response across multiple retinotopic visual field locations. Because there are systematic differences in the cortical folding patterns of different areas, this reduces the ability of neighbouring areas to mimic one another. By using individual MRI, these previous RCSE approaches further leverage individual-specific, finer variations in these cortical folding patterns for a potentially richer set of area-specific constraints. However, this key advantage of the approach is also its main challenge because small undulations in individual cortical surfaces mean that small misspecifications in the retinotopy (due to noisy fMRI measurements) can lead to large variability in dipole orientation, which impacts heavily on forward model estimates. This challenge has been addressed in a number of ways. J. Ales et al (2010) slightly repositioned equivalent current dipoles based on EEG data fit to correct for fMRI misspecification. Hagler and Dale (2013) argued that this gave dipoles too much freedom to vary in orientation and instead opted to model extended cortical patches that would be less sensitive to position misspecification. Because this alone did not eliminate the problem of fMRI noise they implemented an iterative reweighting approach to dampen the influence of visual locations whose residual error stood as outliers. Hagler (2014) developed the approach further by allowing the extended cortical patches to reposition slightly to better match grand average source estimates, yielding more consistent estimates.

In this study, we adopted a distinct grand-average based approach with the main advantage that it eliminates the need for EEG-informed adjustments of model parameters such as generator location and orientation and the weighting of visual field locations in models, reducing flexibility which, in principle, may contribute to cross-talk to some degree. Despite this benefit, there are some drawbacks to the grand-average approach that are important to highlight. The cortical surface of grand-average MRI smooths out the finer details of cortical folding, which may reduce the number of area-specific constraints available. Another issue is that the topography predicted from a grand average brain is not the same as the grand average of topographies generated by individual brains. This is because when topographies are averaged, all the individual topographic variability remains present in the average, producing a diffuse scalp distribution. By contrast, when the cortical surface is averaged first then this variability is removed before the topography is produced and therefore does not feature in it. Thus, the use of grand average MRI retinotopy may misestimate grand average topographies to some extent. However, the consequences of this are unlikely to be prohibitive as it is difficult to see how it would lead to predicted topographies that match VEP topographies artifactually. It is more likely that they would fail to predict some aspects of VEP topographies that would have been predicted by accurately specified individual retinotopy. Therefore, we would contend that this limitation affects the sensitivity but not the specificity of the approach. In relation to the results presented in this study, we therefore suggest that the dominance of V1 in explaining the VEP signals is robust, that the latency separation of V1 from extrastriate areas is robust (if perhaps uncertain in degree), but, as always, that there are no extrastriate contributions at all during the C1 time window cannot be concluded with certainty. Although these limitations could be mitigated to some extent by using individual retinotopy, particularly the second limitation, our choice of approach was ultimately motivated by the goal of the analysis. This was to highlight the topographical features that are uniquely diagnostic of early visual areas so that EEG researchers can make informed source judgements in the absence of fMRI data. In so doing, we identified a heuristic that can distinguish a V1 source from V2/V3 sources based on just two stimulus locations - reversing polarity between the upper and lower visual field along with a tendency towards ipsilateral topographies in the upper field and contralateral ones in the lower visual field. Our results suggest similar heuristics to identify V2 and V3 sources as well. In both cases, signals should reverse in polarity between the upper and lower visual field, as with V1, but in the case of V2, topographies should be slightly contralateral in the upper field and ipsilateral in the lower field, while for V3, topographies should be closer to midline but slightly contralateral at all visual field locations. By highlighting these heuristics, our study allows for certain inferences about visual area sources to be made using EEG alone, broadening the scope of source inferences to a wider range of studies.

## DATA AND CODE AVAILABILITY

Task scripts and MATLAB code for carrying out analyses are available at https://osf.io/gx8vm/?view_only=b014821842564e28812a11a33d72934c.

## AUTHOR CONTRIBUTIONS

Kieran Mohr: Conceptualization, Data curation, Methodology, EEG analysis & forward modelling, Visualisation, Writing - first draft, Writing - review & editing, Funding acquisition, and Project administration.

Anna Geuzebroek: Retinotopy analysis, and Writing - review & editing.

Simon Kelly: Conceptualization, Methodology, Writing - review & editing, Funding acquisition, and Supervision.

## DECLARATION OF COMPLETING INTEREST

The authors declare no competing financial interests.

## ACKNOWLEDGEMENTS

This work was funded by Irish Research Council (GOIPG/2016/123 - to K.S.M.) and The Wellcome Trust (219572/Z/19/Z - to S.P.K.). The funders had no involvement in the study.

## SUPPLEMENTARY MATERIAL

**Table S1.** Latencies of positive (P) and negative (N) peaks of *β*-coefficients in the transient VEP for single-area models.

|  | Polarity (P/N) | Polarity (P/N) | Polarity (P/N) | Polarity (P/N) |
| --- | --- | --- | --- | --- |
|  | Latency (ms) | Latency (ms) | Latency (ms) | Latency (ms) |
| **V1** | P | N | P | P |
|  | 80 | 150 | 195 | 295 |
| **V2** | P | N | N | - |
|  | 105 | 170 | 230 | - |
| **V3** | N | P | N | N |
|  | 75 | 130 | 190 | 305 |
| **MT** | N | P | P | N |
|  | 80 | 155 | 230 | 285 |

**Table S2.** Latencies of peaks and troughs of *β*-coefficients in the transient VEP for the full model (V1-3+MT)

|  | Polarity (P/N) | Polarity (P/N) | Polarity (P/N) | Polarity (P/N) |
| --- | --- | --- | --- | --- |
|  | Latency (ms) | Latency (ms) | Latency (ms) | Latency (ms) |
| **V1** | P | N | P | P |
|  | 80 | 145 | 195 | 295 |
| **V2** | N | P | N | N |
|  | 70 | 115 | 195 | 290 |
| **V3** | N | P | N | N |
|  | 75 | 135 | 220 | 305 |
| **MT** | N | P | P | N |
|  | 80 | 150 | 230 | 285 |

**Table S3.** Latencies of peaks and troughs of *β*-coefficients in the multifocal VEP for single-area models.

|  | Polarity (P/N) | Polarity (P/N) | Polarity (P/N) |
| --- | --- | --- | --- |
|  | Latency (ms) | Latency (ms) | Latency (ms) |
| **V1** | P | N | P |
|  | 75 | 140 | 260 |
| **V2** | P | N | P |
|  | 95 | 145 | 240 |
| **V3** | N | N | N |
|  | 75 | 160 | 310 |
| **MT** | N | P | N |
|  | 85 | 140 | 260 |

**Table S4.** Latencies of peaks and troughs of *β*-coefficients in the multifocal VEP for the full model (V1-3+MT)

|  | Polarity (P/N) | Polarity (P/N) | Polarity (P/N) |
| --- | --- | --- | --- |
|  | Latency (ms) | Latency (ms) | Latency (ms) |
| **V1** | P | N | P |
|  | 75 | 140 | 260 |
| **V2** | N | N | - |
|  | 70 | 150 | - |
| **V3** | N | N | - |
|  | 75 | 160 | - |
| **MT** | N | P | N |
|  | 80 | 140 | 260 |

